# Two teosintes made modern maize

**DOI:** 10.1101/2023.01.31.526540

**Authors:** Ning Yang, Yuebin Wang, Xiangguo Liu, Minliang Jin, Miguel Vallebueno-Estrada, Erin Calfee, Lu Chen, Brian P. Dilkes, Songtao Gui, Xingming Fan, Thomas K. Harper, Douglas J. Kennett, Wenqiang Li, Yanli Lu, Jingyun Luo, Sowmya Mambakkam, Mitra Menon, Samantha Snodgrass, Carl Veller, Shenshen Wu, Siying Wu, Yingjie Xiao, Xiaohong Yang, Michelle C. Stitzer, Daniel Runcie, Jianbing Yan, Jeffrey Ross-Ibarra

## Abstract

Despite its global importance as a crop with broad economic, dietary, and cultural importance, the origins of maize and its closest wild relatives remained the topic of vigorous debate for nearly a century. Molecular analyses ultimately concluded that maize was domesticated once from a common ancestor with its closest extant relative, the lowland wild grass *Zea mays* ssp. *parviglumis*. But neither the current genetic model nor earlier models based on archaeological data account for the totality of available data, and recent work has highlighted the potential contribution of a second wild relative, the highland *Zea mays* ssp. *mexicana*. Here we present a detailed population genetic analysis of the contributions of both wild taxa to modern maize diversity using the largest sample of traditional maize varieties sequenced to date. We show that all modern maize can trace its origin to an ancient admixture event between domesticated ancient maize and *Zea mays* ssp. *mexicana* in the highlands of Mexico ca 5300 cal BP, some 4,000 years after domestication began. We show that variation in admixture is a key component of modern maize genetic and phenotypic diversity, both at the level of individual loci and as a factor driving a substantial component of additive genetic variation across a number of agronomic traits. Our results clarify the long-debated origin of modern maize, highlight the potential contributions of crop wild relatives to agronomic improvement, and raise new questions about the anthropogenic mechanisms underlying multiple waves of dispersal throughout the Americas.

**One-Sentence Summary:** Our results clarify the long-debated origin of modern maize and highlight the contributions of crop wild relatives to the agronomic improvement of modern varieties.

## Main text

The domestication of crops transformed human culture. For many crops, the wild plants that domesticates are most closely related to can be readily identified by morphological and genetic similarities. But the origins of maize (*Zea mays* subsp. *mays* L.) have long been fraught with controversy, even with its global agricultural importance, ubiquity, and extended scrutiny as a genetic model organism. While there was general agreement that maize was most morphologically allied to North American grasses in the subtribe Tripsacinae (*1*,*2*), none of these grasses bear reproductive structures similar to the maize ear, in which seeds are exposed along a compact, non-shattering rachis. The form is so radically distinct from its relatives that the maize ear has been called “teratological” (*3*) and a “monstrosity” (*4*).

Explanations for the ancestry of maize have long been contentious (*5*). A popular model, based on extensive evaluation of the morphology of archaeological samples, argued that modern maize was the result of hybridization between an ancestral wild maize and another wild grass (*6*). This archaeological model, however, fails to explain cytological (*7*) or genetic (*8*,*9*) data showing that maize is most closely related to the extant teosinte *Zea mays* ssp. *parviglumis* (hereafter *parviglumis*). Today, the most widely accepted model is also the simplest – maize was domesticated from a wild annual grass in the genus *Zea*, commonly known as teosinte. This idea, originating with Ascherson (*10*) and championed by George Beadle throughout the 20^th^ century (*4*,*7*), became firmly cemented in the literature after genetic analysis revealed clear similarities between maize and teosinte (*8*,*9*,*11*). Nonetheless, this simple genetic model is insufficient to explain disparities between genetic and geographic overlap between maize and *parviglumis* (*12*)or morphological support for admixture in archaeological samples (*13*–*15*).

Much of the early work on maize origins was complicated by the relatively poor characterization of the diversity of annual teosinte (*16*). In addition to the lowland *parviglumis*,the other widespread annual teosinte is *Z. mays* ssp*. mexicana* (hereafter *mexicana*), found throughout the highlands of Mexico. These taxa diverged 30-60,000 years ago (*17*,*18*) and show clear morphological (*19*), ecogeographic (*20*,*21*), and genetic (*22*,*23*) differences as well as strong evidence for local adaptation along elevation (*24*). In contrast to the overall genetic similarities between maize and *parviglumis*, some early genetic studies identified significant sharing of alleles between *mexicana* and highland maize (*25*), a result confirmed by extensive genome-wide data (*26*,*27*). Maize and *mexicana* co-occur in the highlands of Mexico, but recent work has revealed *mexicana* ancestry far outside this range, including in ancient maize from New Mexico (*28*), modern samples in the Peruvian Andes (*29*), and individual alleles apparently selected broadly in modern maize (*30*,*31*).

### *mexicana* admixture is ubiquitous in modern maize

Archaeological data suggest that after its initial domestication in the lowlands of the Balsas River basin, maize was introduced to the highlands of central Mexico ~6,200 cal BP (calendar years before present)(*32*), where it first came into sympatry with *mexicana*. By this time, however, maize had already reached Panama (by ~7,800 cal BP) (*33*), and even farther into S. America (by ~6,900-6,700 cal BP) (*34*–*36*). Samples from S. America that reflect dispersal events prior to maize colonization of the Mexican highlands should not exhibit evidence of admixture with *mexicana*.Indeed, tests of admixture find no evidence of *mexicana* ancestry in N16, a ~5,500 cal BP maize cob from northern Peru (*37*).

To investigate evidence of *mexicana* admixture across a broad sampling of maize, we applied f4 tests (*38*) using a sample of the diploid perennial teosinte *Zea diploperennis* (*39*) as the outgroup. The greatest diversity of maize is found in present-day Mexico, but whole genome resequencing exists for only a handful of traditional Mexican maize (*40*). We therefore sequenced 267 accessions of open-pollinated traditional maize from across Mexico (**Fig. 1a, table S1 and S2**). Applying f4 tests revealed significant non-zero admixture with *mexicana* in all maize except the ancient Peruvian sample N16 (**Fig. 1b, table S3**). We find evidence for *mexicana* admixture well outside of Mexico, including in modern samples from the Southwest US and the Andes (*40*) and a newly sequenced set of 73 traditional Chinese varieties that post-date European colonization of the Americas (**table S1 and S2**). We extended our search to ancient samples, again finding *mexicana* admixture in archaeological samples from the Tehuacan Valley in central Mexico dating to ~5,300 cal BP (*41*) and both lowland and highland (>2000masl) samples from S. America dating to ~1,000 cal BP (*42*). Finally, we turned to modern breeding material, where again f4 tests identify significant admixture in a diversity panel of more than 500 modern inbred lines (*43*). In sum, we find evidence of *mexicana* ancestry in all examined maize samples dating as early as ~5,300 cal BP.

**Fig. 1.**
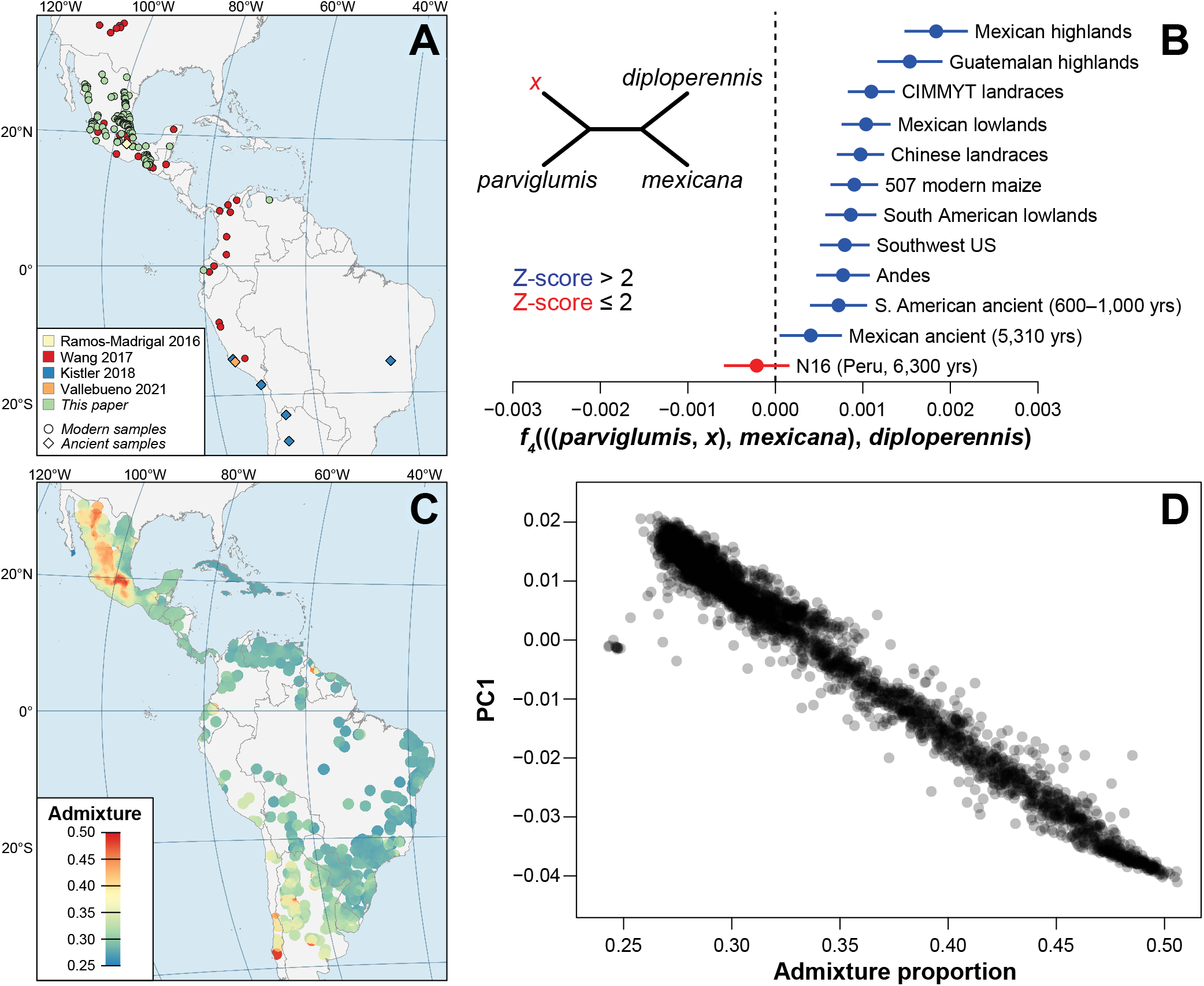
Admixture from *Zea mays* ssp. *mexicana* is ubiquitous in maize. (**A**) Sampling of newly sequenced, published, and ancient maize genomes. See table S1 for details on sampling. (**B**) f4 statistics for different groups of maize. (**C**) Proportion of *mexicana* admixture estimated for ~5,000 field collections from CIMMYT. (**D**) Correlation (r=0.97) between the first principal component of genetic diversity in 5,684 CIMMYT maize landraces and *mexicana* admixture.

To more broadly investigate the importance of introgression to maize diversity, we ran STRUCTURE (*44*) to estimate *mexicana* ancestry in genotyping data from a much larger sample of 5,684 traditional maize varieties and hundreds of wild samples of both subspecies from across the Americas (*45*,*46*). These maize samples also show ubiquitous evidence of *mexicana* admixture (**Fig. 1c**). More surprisingly, principal component analyses of these maize samples reveals that the major axis of genetic variation across maize in the Americas is nearly perfectly correlated with *mexicana* admixture (R^2^=0.97; **Fig. 1d**), suggesting a prominent role for variation in *mexicana* ancestry in patterning genetic diversity in contemporary maize.

### A novel model of modern maize origins

These findings of admixture suggest a pivotal role for *mexicana* in the ancestry of modern maize. To better understand the extent and timing of *mexicana* admixture in maize evolution, we fit a series of admixture graphs to the *f4* statistics for various groups of maize (**fig S1-11**). The resulting graph (**Fig. 2A**) shows an initial domestication from *parviglumis* followed by a deep split between extant maize and the ancient S. American sample N16, dating to ~5,500 cal BP. Subsequent to this split, the dispersal and adoption of maize by people living in the highlands of Mexico led to significant admixture with *mexicana*. An independent genetic estimate of the timing of this admixture (**Supplementary Materials**) resulted in high uncertainty, but the point estimate of 5,716 yrs (±5,614) is consistent with the earliest archaeological evidence of maize in the Mexican highlands (~6,200 cal BP) (*47*).

**Fig. 2.**
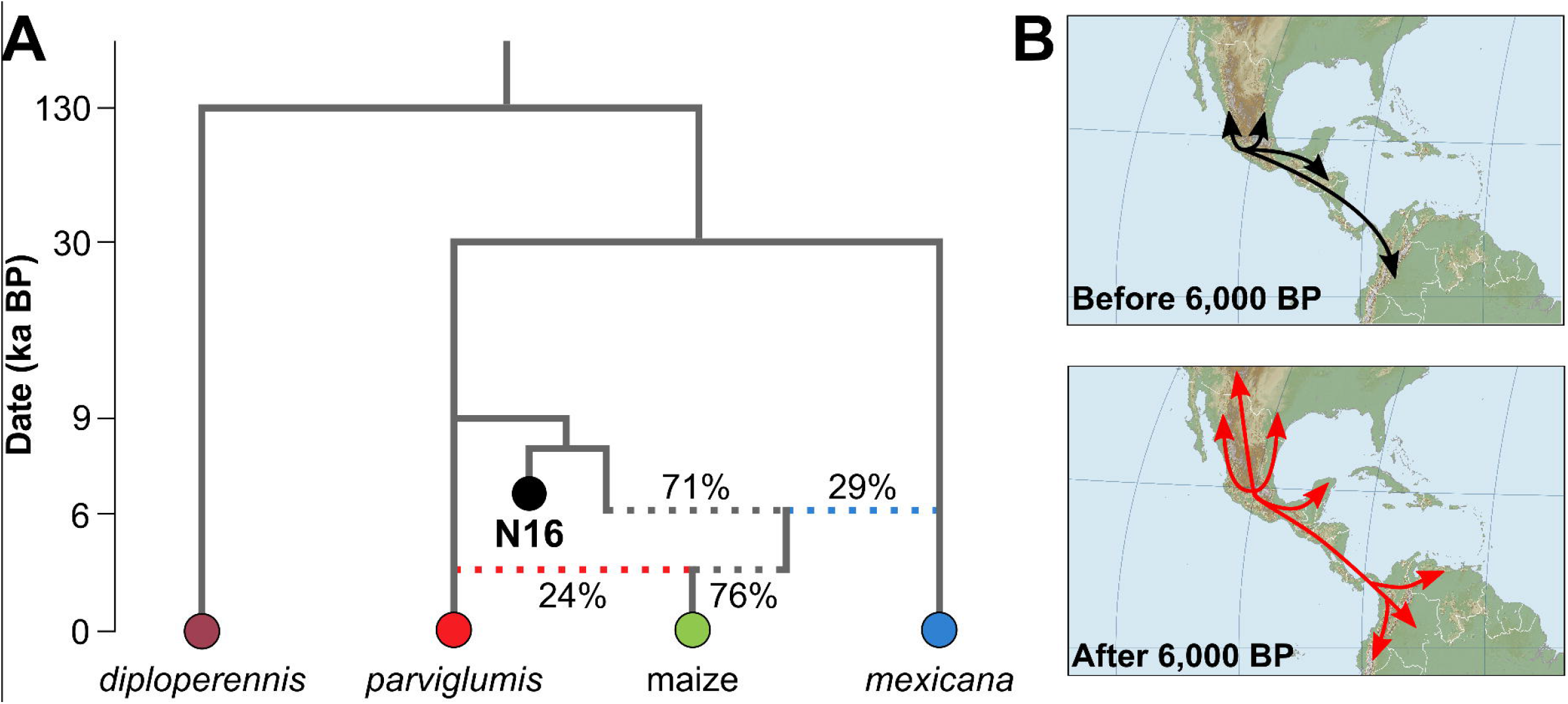
A novel model of maize origins. (**A**) Admixture graph showing two admixture events (dotted lines) in the history of modern maize. Shown are the estimated percentage contributions of each donor population for both hybridization events. Extant maize is represented by a diverse panel of ~500 inbred lines (**Supplementary Materials**). (**B**) Proposed model of maize origin showing two waves of movement out of Mexico: early movement after initial domestication in the Balsas (top; black) and a second wave out of the highlands of Mexico after admixture with *mexicana* (bottom; red).

*mexicana* ancestry proportions in maize range from 21.7% to 34.7% in the fitted admixture graphs, with the highest values estimated for traditional varieties from the highlands of Mexico and Guatemala, consistent with initial f4 statistics (**Fig. 1B**). Our graph-based estimate for a broad panel of more than 500 diverse modern inbreds estimates an average of 22% admixture (**fig. S1**), highlighting the important contribution of *mexicana* to modern maize. While these estimates are lower than those from reduced representation genotyping (**Fig. 1C**), genotyping SNPs overestimate genome-wide admixture proportions because of their biased distribution across the genome (**Supplementary Materials, fig. S12**).

We interpret the ubiquitous admixture we observe as evidence supporting a new model of maize origins (**Fig. 2B**). Consistent with previous work (*42*,*48*), we propose that maize dispersed out of the Balsas River basin in Mexico, quickly reaching S. America by at least ~6,500 cal BP (*49*). Then, ~6,000 cal BP, maize was adopted by people living in the highlands of central Mexico where it admixed with *mexicana* (*12*,*26*,*27*). Our data suggest that this admixed maize then spread widely from the highlands of Mexico, replacing or mixing with existing populations across the Americas, introducing *mexicana* alleles as it moved. As it moved into the lowlands of Mexico, maize once again came into contact with *parviglumis*, and our admixture graph suggests this secondary contact resulted in substantial additional *parviglumis* ancestry. This model is consistent with the second wave of maize admixture into S. America posited by Kistler *et al*. (*50*), but further explains the origin of that wave and the existence of *mexicana* alleles far outside *mexicana’s* native range in S. America by at least ~1,000 cal BP. The two-admixture graph fits f4 statistics for all extant and ancient maize samples (**fig. S1-11**), and simpler graphs that omit one or two of these admixture events do not fit the data well (**fig. S13-15**). Finally, the existence of the N16 maize cob dating to 5,500 cal BP – which shows no *mexicana* admixture – allows us to distinguish this model from one in which maize was domesticated from admixed populations of teosinte in one of the hybrid zones between *mexicana* and *parviglumis* (**fig. S16-S17**) (*51*).

The timing of admixture between maize and *mexicana* admixture in the highlands of Mexico between 6,000 and 4,000 cal BP corresponds with observed increases in cob size and number of seed rows from archaeological samples (*52*,*53*). Southward dispersal of maize varieties with *mexicana* admixture parallels a northward flow of people (~5,600 cal BP) originating as far south as Costa Rica and Colombia, and coincides with the appearance of improved maize varieties in Belize (*50*,*54*). Archaeological samples demonstrate the presence of maize as a staple grain in the neotropical lowlands of Central America subsequent to *mexicana* admixture, between 4,700 and 4,000 cal BP (*55*,*56*). Ultimately, all varieties of maize in Mesoamerica had *mexicana* admixture by ~3,000 cal BP as it became a staple grain across the entire region (*28*,*53*,*56*,*57*).Early Mesoamerican sedentary agricultural villages developed at this time along with the appearance of hereditary leadership evident in later state-level societies dependent upon more intensive forms of maize agriculture (*58*–*60*). The confluence of archaeological and genetic data thus suggests that *mexicana* admixture was central to the widespread use and dispersal of maize in the Americas after ~4,000 cal BP.

### Variation in admixture along the genome

Having established a central role for both *parviglumis* and *mexicana* in the origins of modern maize diversity, we next explored variation in *mexicana* ancestry across the genome. Using un-admixed *parviglumis* and *mexicana* individuals (*39*) as references, we applied an ancestry hidden Markov model to identify regions of *mexicana* ancestry along individual maize genomes (see Methods). In close agreement with our admixture graph, we estimate 15-25% average *mexicana* ancestry across 845 maize genomes (mean 18%; **table S4**) (*61*). This variation in total ancestry among modern maize is much greater than that predicted from a single pulse of ancient admixture (**Supplementary Materials**), and likely reflects a combination of selection as well as ongoing gene flow in parts of the range (*27*). Mean *mexicana* ancestry also varies considerably along the genome (**Fig. 3, A and B**), though 80% of introgression tracts are smaller than 9,113 bp (**fig. S18**), consistent with a relatively ancient origin of most *mexicana* admixture.

**Fig. 3.**
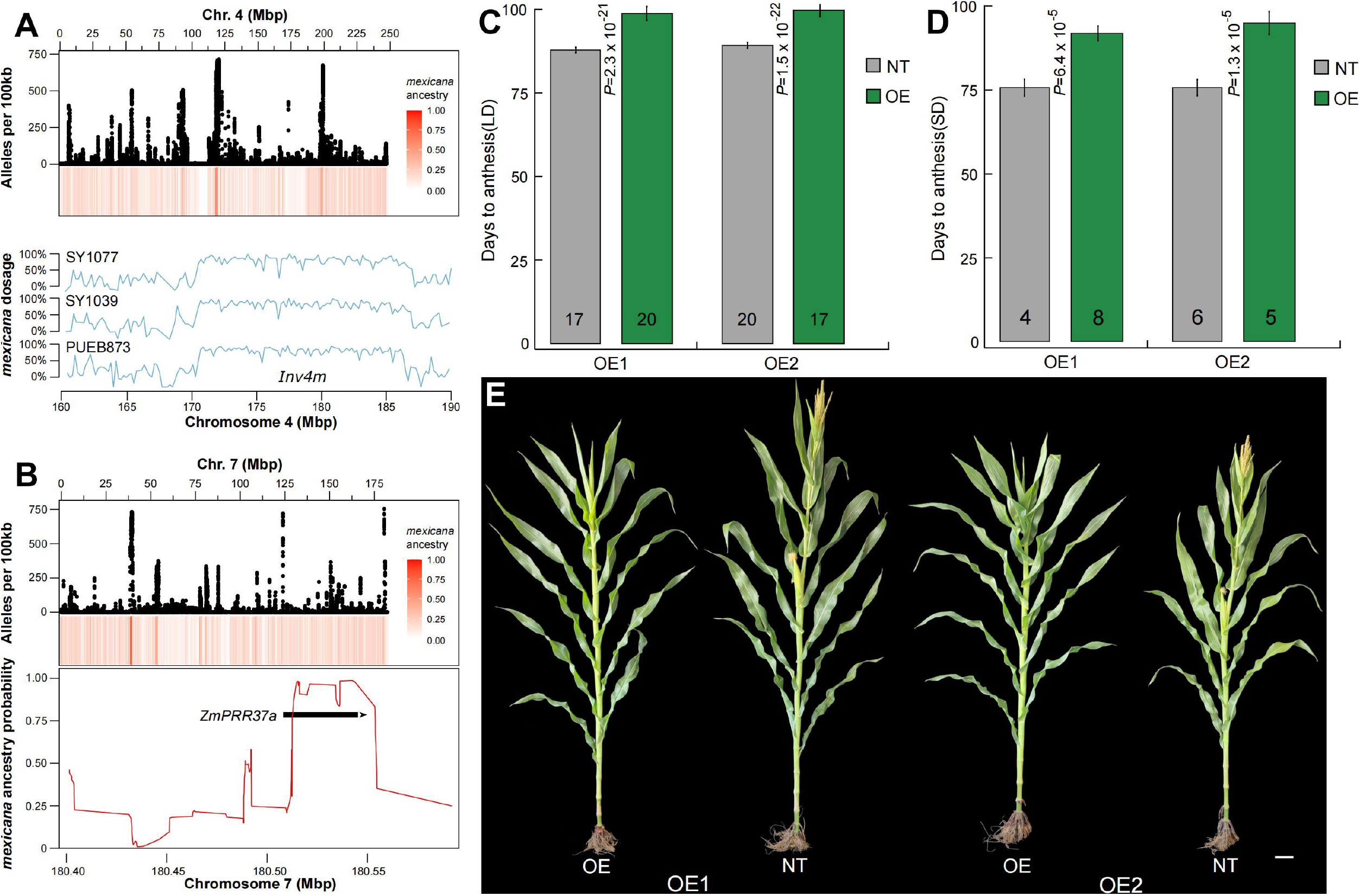
Variation and functional validation of *mexicana* admixture. (**A**) (top) number of high frequency (>80%) *mexicana* alleles along chromosome 4 (black points) and average *mexicana* ancestry (red). (bottom) *mexicana* ancestry of 3 inbred lines in the region around chromosome inversion *Inv4m*. (**B**) (top) number of high frequency (>80%) *mexicana* alleles along chromosome 7 (black points) and average *mexicana* ancestry (red). (bottom) *mexicana* ancestry in B73 across the *ZmPRR37a* gene model (black bar). The differences of days to anthesis for nontransgenic (NT) and overexpression (OE) lines of *ZmPRR37a* in (**C**) long day conditions (2022, China, E124°49’, N43°30’) and (**D**) shortday conditions(2021, China, E108°43’, N18°34’). The data in (**C and D**) are means ±SE. The numbers in each column indicate the sample sizes. The level of significance was determined using one-way analysis of variance(ANOVA). (**E**) Nontransgenic (NT) and two independent overexpression (OE) lines of *ZmPRR37a* grown in long day conditions. Scale bar=10 cm.

In addition to the majority of small tracts, however, we also identify numerous signals consistent with an important role for inversion polymorphisms. These include the apparent presence of the large inversion *Inv4m* – a well-studied target of adaptive introgression in maize from highland environments (*26*,*27*) – in two Chinese inbred lines and one traditional Mexican variety (**Fig. 3A**). We also see high levels of *mexicana* admixture in the region of *Inv1n*, a 50Mb inversion common in *parviglumis* but rare in *mexicana* and entirely absent in maize (*63*) (**fig. S19**). Finally, we estimate dramatically decreased admixture for chromosomes 8 and 9 (**fig. S20**), which we hypothesize is due to the presence of multiple large *mexicana*-specific inversions which could hinder introgression by repressing recombination (*18*,*22*).

A detailed look at admixture along individual genomes also enabled us to begin to investigate the functional significance of *mexicana* admixture. We identified regions of the genome in which high confidence *mexicana* alleles (>90% posterior probability) were at high frequency (>80%) across all modern maize (see Methods), consistent with recent positive selection (**Fig. 3, A and B**). We found these loci clustered into eleven regions, which overlap QTL for agronomically relevant phenotypes (*64*) and include a number of genes with well-studied functions in *Arabidopsis* such as disease resistance and floral morphology (**table S5**). We focused on one region on chromosome 7, where we found a narrow peak of high frequency *mexicana* alleles that overlaps with a maize-teosinte flowering time QTL(*64*) and is centered on the gene *Zm00001d022590*, also known as *ZmPRR37a* (**Fig. 3B**). *Mexicana* alleles at *ZmPRR37a* SNPs are found in up to 89% of all maize, including the reference genome line B73 (**fig. S21**). *ZmPRR37a* is thought to be involved in the circadian clock-controlled flowering pathway (*65*) and is an ortholog of the sorghum gene *Ma1* which controls flowering under long-day conditions (*66*). To validate this function, we obtained a CRISPR/Cas9 knockout mutant from a targeted mutagenesis library (*67*) and also developed two transgenic overexpression lines (**Supplementary Materials**). Consistent with its hypothesized role in response to daylength, *ZmPRR37a* knockout mutants exhibited significantly earlier flowering phenotype in long day conditions (**fig. S22, A, B and D**) but show no effect in short day conditions (**fig. S22, A and C**), and overexpression lines exhibited significantly late flowering in both long and short day conditions (**Fig. 3, C-E**). Maize carrying the *mexicana* introgression at *ZmPRR37a* show lower levels of expression than *parviglumis* (*68*), and our functional evaluation thus suggests *mexicana* alleles at *ZmPRR37a* may have helped maize adapt to earlier flowering in long-day conditions as it expanded out of Mexico to higher latitudes.

### *mexicana* admixture underlies phenotypic variation in maize

Admixture with teosinte has been associated with phenotypic variation for a number of traits in traditional maize (*69*). Our analysis of *parviglumis* ancestry replicates these historical findings (**fig. S23, table S6**), and *mexicana* gene flow has been instrumental in the phenotypic adaptation of maize to the highlands (*26*, *70*–*72*). If *mexicana* admixture played a key role in the dispersal and use of maize, *mexicana* alleles should contribute to agronomically relevant phenotypic variation. We thus combined our estimates of admixture with data from 33 phenotypes to perform multivariate admixture mapping across 452 maize inbreds (**Supplementary Materials**). We find 92 associations at a false discovery rate of 10%. Grouping these into 22 peaks, we identify 25 candidate genes within 5kb of the peak SNP (**Fig. 4, figure S24, table S7 and S8**). These include a significant association with zeaxanthin – a carotenoid pigment that plays a role in light sensing and chloroplast movement (*73*) and is of significance to human health (*74*) – approximately 1kb downstream of the gene *ZmZEP1*, a key locus in the xanthophyll cycle that regulates zeaxanthin abundance in low light conditions (**fig. S25A**). Haplotype visualization reveals clear sharing between maize and *mexicana* (**fig. S25B**), and the *mexicana*-like haplotype increases the expression of *ZmZEP1* and reduces zeaxanthin content in maize kernels (**fig. S25C**). *Mexicana* ancestry at *atm1* is associated with changes in kernel width, and previous work has identified how variation at *atm1*, in conjunction with the maize gene *atr*, can impact kernel size and starch content (*75*). We also see associations with well-known lipid metabolism genes such as *dgat1* and *fae2* (*76*). The *mexicana* allele at *dgat1* is associated with a decrease in the proportion of linoleic acid but an increase in overall oil content, and is independent of the well-studied amino acid variant (*77*). Though expression of dgat1 has been suggested to play a role in cold tolerance in maize and *Arabidopsis* (*78*,*79*), a preliminary experiment in maize seedlings failed to identify differences in cold tolerance in lines of varying ancestry at *dgat1* (**fig. S26**). In addition to compelling candidate loci in modern inbreds, we applied a novel genotype by environment association mapping approach (*80*) in a large set of maize landraces (*81*). We find a strong novel association on chromosome 1 (**Fig. 4**), where *mexicana* ancestry increases cob size. The candidate gene closest to the associated SNP, *Zm00001d029675*, was recently identified as a target of selection during breeding efforts in both the U.S. and China (*82*).

**Fig. 4.**
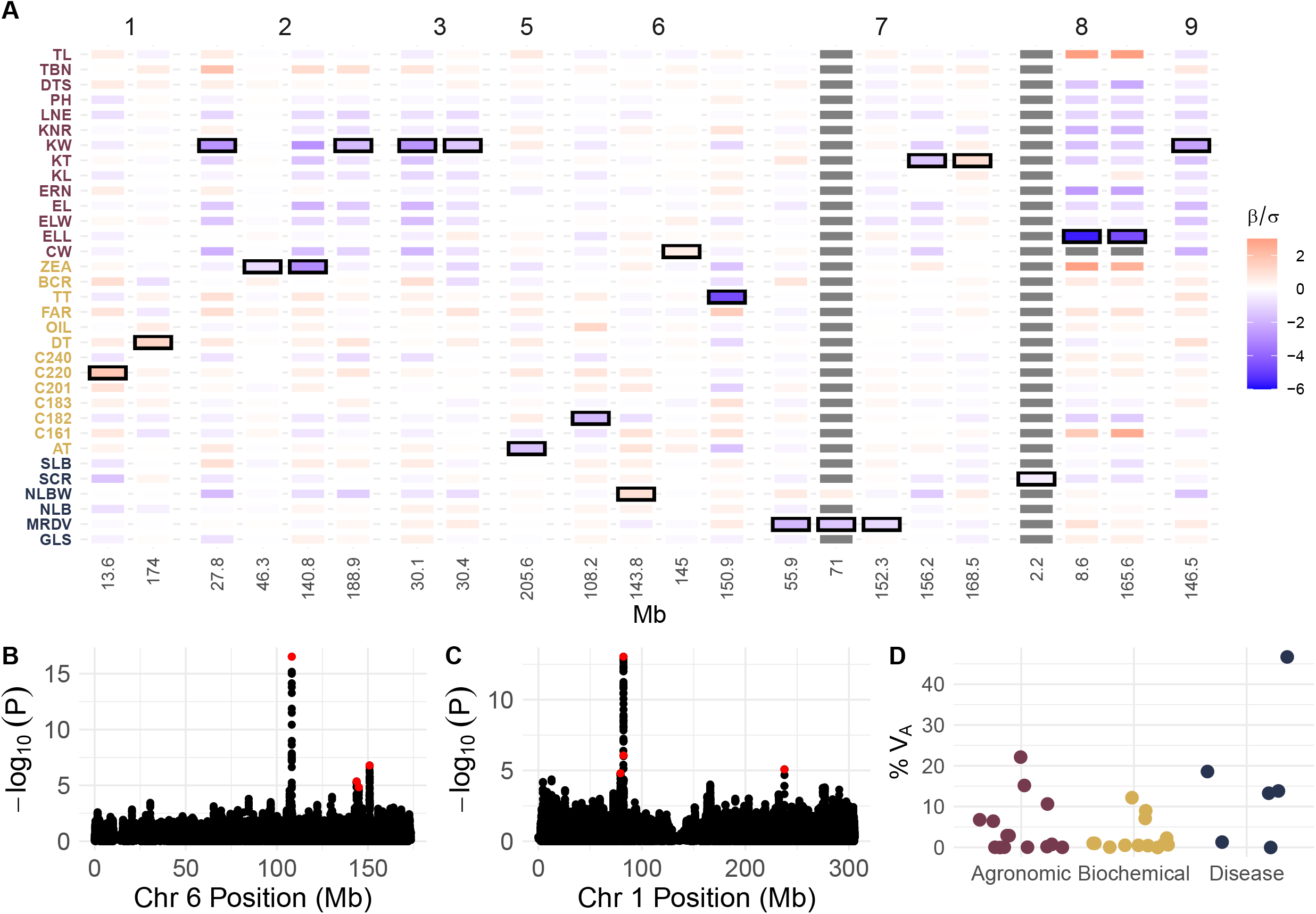
Phenotypic impacts of *mexicana* admixture. (**A**) Effect sizes (scaled by trait standard deviation) across traits for the 22 lead SNPs from admixture GWAS in the inbred diversity panel. Grey boxes represent missing data due to low minor allele frequency. Black outlines show the trait with the largest absolute value effect size for each SNP. Numbers above each group of columns represent chromosomes, while numbers below represent Mb positions. Trait name acronyms and description are in **table S9**, trait acronym colors represent categories shown in (**D**). (**B**) Manhattan plot of admixture GWAS for linoleic acid content in the inbred diversity panel. The peak includes the gene *dgat1*. (**C**) Manhattan plot of admixture GWAS for cob weight using traditional maize varieties. Red points in (**B**) and (**C**) represent the lowest p-value SNP within 500kb windows around significant associations. (**D**) Variance partitioning in the inbred diversity panel. Shown is the proportion of additive genetic variance explained by *mexicana* admixture.

While GWAS approaches can identify individual loci with large effects, it is likely that *mexicana* admixture contributes important variation of smaller effect size to a number of polygenic traits. To test this hypothesis, we used the same inbred association panel to estimate the proportion of additive genetic variance contributed by *mexicana* across a set of 33 phenotypes (**Supplementary Materials**, **table S9**). We estimate that *mexicana* admixture explains a meaningful proportion of the additive genetic variation for many of these traits, including nearly 25% for the number of kernels per row, 15% for plant height, 10% for flowering time, approximately 5-10% of several disease phenotypes, and 15 to nearly 50% for multiple disease phenotypes (**Fig. 4D**).

## Discussions

Conflicting archaeological, cytological, genetic, and geographic evidence led to two irreconcilable models for the origin of maize. Here, using the broadest sampling to date of whole genome sequence of traditionally cultivated maize and teosinte, we revisited the evidence for admixture between maize and its wild relative *Zea mays* ssp*. mexicana*. We propose a new model which posits that, after admixture with *mexicana* in the highlands of central Mexico, admixed maize spread across the Americas, either replacing or hybridizing with pre-existing maize populations. While this model is consistent with both genetic and archaeological data, it nonetheless raises a number of questions. Among these, most notable is perhaps the question of why and how this secondary spread occurred - was it due to some advantage of the admixed maize over earlier domesticated forms, or was the spread coincidental with demic or cultural exchange among human populations (*54*)?

Changes in maize cob morphology and dietary isotope data from human populations in Central America indicate a transition between early cultivation and the use of maize as a staple grain between 4,700 and 4,000 yr B.P. (*55*). This timing suggests a possible direct role for hybridization between maize and *mexicana* in improving early domesticated forms of maize. To better understand why admixed maize may have been beneficial for early farmers, we sought to investigate phenotypic associations between *mexicana* alleles and phenotypes in extant maize. We identify and functionally validate a locus important for photoperiodicity and flowering time, and find candidate genes associated with important agronomic phenotypes including nutritional content and kernel and cob size. None of these loci individually, however, are likely sufficient to drive a large advantage of admixed maize. And while we show that, combined, alleles introgressed from *mexicana* explain a meaningful proportion of additive genetic variance for agronomic and disease resistance traits, it remains unclear whether this novel variation could drive rapid adoption of admixed maize. We speculate that, in addition to variation at these specific phenotypes, admixture may have played a role in the spread of admixed maize by augmenting genetic diversity and ameliorating genetic load in early domesticated populations, perhaps even providing some generalized hybrid vigor. Indeed, the global ecological niche of cultivated maize more closely reflects that of *mexicana* than *parviglumis* (*61*), and *maize-mexicana* hybrids show extensive heterosis for both viability and fecundity. Modern ethnographic evidence is also consistent with these ideas, as farmers continue to introgress teosinte into their maize populations to make them “stronger” (*16*,*83*,*84*).

Introgression between relatives has long been recognized as a major source of plant adaptation (*85*), yet only with the advent of molecular markers have we begun to recognize the key role that gene flow from wild relatives has played in crop evolution (*86*). Here, with unprecedented sampling and genomic coverage of both traditional and modern varieties as well as wild relatives and ancient samples, we argue that introgression from a close wild relative of maize was pivotal to its success as a staple crop. The presence of adaptive variation in wild relatives is not unique to maize, and we predict a similar history will be revealed for many other crops. Indeed, preliminary results already suggest key roles for hybridization in the evolution of rice, tomato, and barley, among others (*87*–*89*). These results not only highlight the past importance of crop wild relatives, but point to their potential as a source of adaptive diversity for future breeding. Most importantly, the work presented here suggests that, for many crops, millenia of diligent efforts by early farmers has capitalized on this diversity and an abundance of relevant functional diversity may already be segregating in traditional varieties or preserved *ex situ* in germplasm gene banks.

## Acknowledgements

This research was supported by funding from the National Natural Science Foundation of China (2022YFD1201500,2020YFE0202300), the National Natural Science Foundation of China (U1901201), Science and Technology major program of Hubei Province (2021ABA011) and the 111 Project Crop Genomics and Molecular Breeding (B20051) to J.Y., the National Natural Science Foundation of China (32222062) to N.Y., the US National Science Foundation (1822330) and US Dept. of Agriculture (Hatch project CA-D-PLS-2066-H 548) to J.R-I., and US National Science Foundation (1546719) to J.R-I. and D.E.R. Most computation resources were provided by the high-throughput computing platform of the National Key Laboratory of Crop Genetic Improvement at Huazhong Agricultural University and supported by H. Liu. We are grateful to R Rellán-’Álvarez, RJ Salvador, RJ Sawers and J Bernal, who provided comments on an early version of this manuscript. Finally, we would like to thank Felix Andrews for statistical advice, although we did not follow it.

## Supplementary Materials

Materials and Methods

Figs. S1 to S30

Tables S1 to S11

