## Supplemental Materials for "Two teosintes made modern maize"

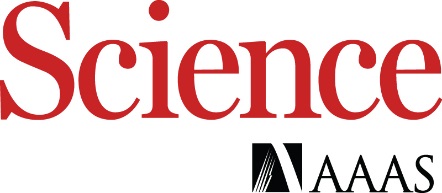

Supplementary Materials for

**Two teosintes made modern maize**

Ning Yang^1,2,3†*^, Yuebin Wang^1,2†^, Xiangguo Liu^4†^, Minliang Jin ^1^, Miguel Vallebueno-Estrada^5^, Erin Calfee^3,6^, Lu Chen^1^, Brian P. Dilkes^7^, Songtao Gui^1^,Xingming Fan^8^, Thomas K. Harper^9^, Douglas J. Kennett^10^, Wenqiang Li^1,2^, Yanli Lu^11^, Jingyun Luo^1,2^, Sowmya Mambakkam^3^, Mitra Menon^3,6^, Samantha Snodgrass^12^, Carl Veller^3,6^, Shenshen Wu^1^, Siying Wu^1^, Yingjie Xiao^1,2^, Xiaohong Yang^13^, Michelle C. Stitzer^14^, Daniel Runcie^15^, Jianbing Yan^1,2*^, Jeffrey Ross-Ibarra^3,6,16*^

^†^These authors contributed equally to this work.

**Materials and Methods**

Samples, whole genome resequencing, reads mapping and SNP calling

SNP data from 507 modern maize inbred lines, 90 *Z.mays* ssp. *mexicana*, 75 *Z. mays* ssp.*parviglumis*, 2 *Z.diploperenis* and 1 *Tripsacum dactyloides* were obtained from version 1 of the ZEAMAP project (*39*,*90*). We sequenced an additional 340 traditional maize varieties, including 267 from across Mexico and 73 from China (**table S1 and S2**). Seeds of each variety were sown in 2018 in Sanya, Hainan Province, China (18°14'35.02"N, 109°30'18"E). Young leaves were collected and frozen at -80°C for DNA extraction. All shotgun libraries were constructed following a standard protocol (*91*) and 150-bp paired-end DNA sequencing on the BGISEQ-500 platform was conducted by BGI (Shenzhen, China). All raw reads are available on NCBI under project PRJNA783885. We followed previous methods (*39*) for read mapping and SNP calling. Raw reads were first processed using FastQC v0.11.3 (*92*) and Trimmomatic v0.33 (*93*) to remove poor-quality base calls and adaptors. Reads were then aligned to the B73 reference genome v4 (*94*) using Bowtie2 v2.1.0, --very-fast (*95*). Unique mapped reads were sorted and indexed using Picard v1.119 (*96*). SAMtools v1.3.1 (*97*) and UnifiedGenotyper from GATK v3.5 (*98*) were used to create the variant calling file for each individual. Hard filtering of individual SNP calls was carried out with mapping quality (MQ ≤ 20.0), and thresholds set by sequencing coverage based on minimum coverage (DP ≤ 5) and maximum coverage (DP ≥ 200). Then, variants from the ZEAMAP and newly sequenced varieties were combined by GATK CombineVariants to a single variant calling file. Newly identified SNPs were recalled in previous samples and replaced with the reference allele if there were supporting reads. Sites with a missing rate higher than 75% across all samples were excluded. The total number of SNPs in ZEAMAP increased from 65,717,599 to 88,149,885. We also collected DNA sequencing data of 30 published traditional varieties and ten ancient maize samples (**table S1**) and called the known sites from the enlarged ZEAMAP of these lines. For all ancient maize samples, we removed all G (reference allele) to A (alternative allele) and C (reference allele) to T (alternative allele) SNPs to avoid the confounding bias caused by the apparent G to A and C to T substitutions in ancient samples (*99*).

*f4* tests and admixture graph

To understand the relationship between ancient maize, modern maize and teosinte populations, we applied f4 statistics analyses (*100*) in admixr (*101*) and built an admixture graph in admixturegraph (*102*). For these analyses, we first identified *parviglumis* and *mexicana* individuals that were not admixed with each other or with maize using the software ADMIXTURE (*103*) and assuming a set of simple K=2 models (**fig. S13**). *Tripsacum dactyloides* was used to polarize alleles as derived/ancestral. We then estimated *f4* (*parviglumis*, *diploperennis*; test population, *mexicana*). We used *Zea diploperennis* as the outgroup and our unadmixed *mexicana* and *parviglumis* as the two contributors to the admixed test population.

Following previous phylogenetic work (*18*), we construct an admixture graph that includes a split between the species *Zea mays* and *Zea* *diploperennis,* followed by a split between *mexicana* and *parviglumis*. Maize is modeled as deriving initially from *parviglumis*, consistent with archaeological (*104*) and genetic (*11*,*12*) data. Paleoecological evidence of slash and burn farming, including microbotanical remains (*58*,*105-113*), support a rapid spread of maize through Central America, appearing in Panama after 8700 cal BP and in the Amazon Basin by 6,900 cal BP (*35*,*36*). Genetic analyses support these results, identifying a deep split between South American and Mesoamerican maize (*42*). We modeled this by adding a split between our ancient Peruvian sample N16 and other ancient and extant maize. Note that results here and in Vallebueno *et al*. (*37*) show no evidence of admixture between *mexicana* and N16. After this initial split, we add an admixture event occurring in the highlands of Mexico (*12*,*26*,*27*) and a subsequent admixture with *parviglumis* as suggested by (*42*). We model N16 as having no subsequent admixture with *mexicana* or *parviglumis* (**fig. S1**). We used the fitting procedure in admixturegraph for fitting graph parameters to *f4* statistics. Bat Cave and McUen samples from de Fonseca *et al*. (*28*) had too few derived alleles to perform *f4*.

Dating introgression

We used the software DATES (Distribution of Ancestry Tracts of Evolutionary Signals) (*113*) to estimate the timing of admixture between *mexicana* to maize. We used the genotypes of a diverse set of 507 maize inbred lines (*43*) as input to DATEs, so DATES estimated the time scale for the whole population. DATES was run with parameters binsize 0.00005, maxdis 1.0, runmode 1, chithresh 0, mincount 1, qbin 50, lovalfit 0.1, and with runfit, afffit and zdipcorrmode set to YES.

*mexicana* admixture in CIMMYT SeeD GBS samples

To evaluate the genomic landscape of introgression across the Americas, we took advantage of a large sample of 5,373 traditional maize varieties and 164 *parviglumis* and 146 *mexicana* for which genotyping-by-sequencing (GBS) genotype data was available (*45*,*46*). To estimate admixture in these individuals, we used the site-by-site ancestry assignment probability with linkage information implemented in STRUCTURE v.2.3 (*44*). This approach was implemented at the level of each chromosome using unphased data and setting *mexicana* and *parviglumis* individuals as members of pre-defined parental populations. The admixture linkage disequilibrium model with K=2 was run in STRUCTURE for a burn-in of 5,000 steps and subsequently 5,000 steps were retained for analysis. Ancestry genotype calls were then summarized into 0,1 and 2 *mexicana* alleles per individual per locus basis for downstream analyses. Admixture results for all individuals are available online (*115*).

Admixture estimated from GBS SNPs likely represents an overestimate of the genome-wide admixture proportion. GBS SNPs are biased towards high recombination regions of the genome (mean 1.7cM/Mb compared to the genome-wide average of 0.75cM/Mb), and *mexicana* admixture is positively correlated with recombination in extant populations of maize sympatric with *mexicana* (*27*). Supporting the idea that GBS admixture proportions are overestimates, sub-sampling our resequencing SNPs to match the distribution of reduced-representation SNPs recovers a higher estimate of admixture (**fig. S12**).

ELAI and high frequency introgressed alleles

We investigated genome-wide patterns of introgression of all the 845 maize and landraces using the software ELAI (*116*). We first ran ADMIXTURE (*103*) using 1,665,173 SNPs from 90 *mexicana* and 75 *parviglumis* individuals after removing SNPs with MAF <0.05 from Chen *et al*. (*39*). From the initial ADMIXTURE of those two taxa with k=2, we chose 48 *mexicana* and 43 *parviglumis* individuals that were unadmixed (membership probabilities in their own cluster of >0.99) to compare in a second model with k=2 to 48 and 43 maize which were randomly selected from 507 modern maize. Individuals showing no admixture with maize were then retained. *parviglumis* was further subdivided given the deep split often seen between populations in the Balsas and Jalisco (*22*). Finally we randomly selected 30 unadmixed accessions each of *mexicana* and Balsas *parviglumis* as reference populations for ELAI (**table S10, fig. S27**). We ran ELAI with two upper-layer clusters and two lower-level clusters and 20 EM steps. We set the timing of admixture to 6,000 generations following the results from DATES. Missing SNPs in the tested individual were filtered and SNPs with a missing rate larger than 0.1 or minor allele frequency smaller than 0.05 in both reference panels were also removed. We analyzed each chromosome of each genome individually as the result of admixture between the haplotypes of the two reference populations, *parviglumis* and *mexicana* (**table S10**). Then we integrated the SNPs with ELAI score in any individual to be an introgression map including 845 individuals with 878,439 SNPs (*61*). We identified tracts of introgression as adjacent SNPs all with ELAI scores larger than 1.8 (*mexicana* dosage >0.9). We also calculated the average genome-wide ancestral allele dosage for all individuals (**table S4**). We defined high frequency *mexicana* alleles as those for which more than 80% of the 845 maize lines had ELAI scores >1.8.

Average estimates of admixture from ELAI were lower than those estimated from GBS data using STRUCTURE. To resolve whether this was due to differences in methodology or data, we compared ELAI estimates of admixture for SNPs within 1kb of GBS markers compared to more distant SNPs. We found that SNPs close to GBS markers had higher average ELAI estimates than the rest of the genome (**fig. S12**), consistent observed differences between the two sets of data. This is also consistent with the bias of GBS markers to genic, high-recombination regions of the genome (*117*) and observed negative correlations between recombination rate and recent *mexicana* ancestry (*27*).

Variance in admixture

In a large well-mixed population,
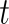
<https://www.codecogs.com/eqnedit.php?latex=t#0> generations after a pulse of admixture, under neutrality, the Morgan length of an introgressed block,
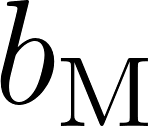
<https://www.codecogs.com/eqnedit.php?latex=b_%5Ctext%7BM%7D#0>, is approximately exponentially distributed with mean
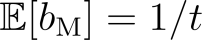
<https://www.codecogs.com/eqnedit.php?latex=%5Cmathbb%7BE%7D%5Bb_%5Ctext%7BM%7D%5D%20%3D%201%2Ft#0> and variance
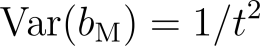
<https://www.codecogs.com/eqnedit.php?latex=%5Ctext%7BVar%7D(b_%5Ctext%7BM%7D)%20%3D1%2Ft%5E2#0> (118). If the overall introgressed fraction is
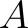
<https://www.codecogs.com/eqnedit.php?latex=A#0>, then the mean number of blocks carried by each individual is
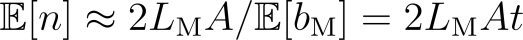
<https://www.codecogs.com/eqnedit.php?latex=%5Cmathbb%7BE%7D%5Bn%5D%20%5Capprox%202L_%5Ctext%7BM%7DA%2F%5Cmathbb%7BE%7D%5Bb_%5Ctext%7BM%7D%5D%20%3D%202L_%5Ctext%7BM%7DAt#0>, where
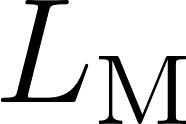
<https://www.codecogs.com/eqnedit.php?latex=L_%5Ctext%7BM%7D#0> is the map length of the genome. If blocks can be assumed to be inherited independently, then the number of blocks an individual carries is approximately Poisson distributed, the variance therefore being
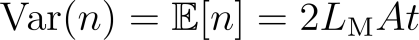
<https://www.codecogs.com/eqnedit.php?latex=%5Ctext%7BVar%7D(n)%20%3D%20%5Cmathbb%7BE%7D%5Bn%5D%20%3D%202L_%5Ctext%7BM%7DAt#0>.

Conditional on receiving
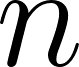
<https://www.codecogs.com/eqnedit.php?latex=n#0> blocks, the fraction of the total Morgan length of an individual's genome that is introgressed,
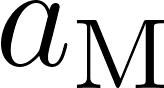
<https://www.codecogs.com/eqnedit.php?latex=a_%5Ctext%7BM%7D#0>, obeys

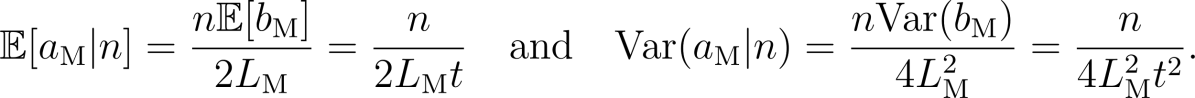
<https://www.codecogs.com/eqnedit.php?latex=%5Cmathbb%7BE%7D%5Ba_%5Ctext%7BM%7D%7Cn%5D%20%3D%20%5Cfrac%7Bn%5Cmathbb%7BE%7D%5Bb_%5Ctext%7BM%7D%5D%7D%7B2L_%5Ctext%7BM%7D%7D%20%3D%20%5Cfrac%7Bn%7D%7B2L_%5Ctext%7BM%7Dt%7D%20%5C%2C%5C%2C%5C%2C%5C%2C%20%5Ctext%7B%20and%20%7D%20%5C%2C%5C%2C%5C%2C%5C%2C%20%5Ctext%7BVar%7D(a_%5Ctext%7BM%7D%7Cn)%20%3D%20%5Cfrac%7Bn%5Ctext%7BVar%7D(b_%5Ctext%7BM%7D)%7D%7B4L_%5Ctext%7BM%7D%5E2%7D%20%3D%20%5Cfrac%7Bn%7D%7B4L_%5Ctext%7BM%7D%5E2t%5E2%7D.#0>

From the law of total variance,

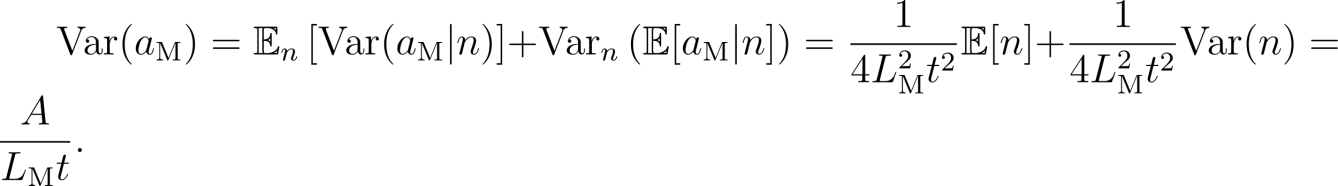
<https://www.codecogs.com/eqnedit.php?latex=%5Ctext%7BVar%7D(a_%5Ctext%7BM%7D)%20%3D%20%5Cmathbb%7BE%7D_n%5Cleft%5B%20%5Ctext%7BVar%7D(a_%5Ctext%7BM%7D%7Cn)%20%5Cright%5D%20%2B%20%5Ctext%7BVar%7D_n%5Cleft(%5Cmathbb%7BE%7D%5Ba_%5Ctext%7BM%7D%7Cn%5D%5Cright)%20%3D%20%5Cfrac%7B1%7D%7B4L_%5Ctext%7BM%7D%5E2t%5E2%7D%5Cmathbb%7BE%7D%5Bn%5D%20%2B%20%5Cfrac%7B1%7D%7B4L_%5Ctext%7BM%7D%5E2t%5E2%7D%5Ctext%7BVar%7D(n)%20%3D%20%5Cfrac%7BA%7D%7BL_%5Ctext%7BM%7Dt%7D.#0>

If
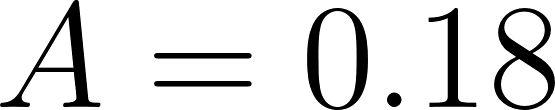
<https://www.codecogs.com/eqnedit.php?latex=A%20%3D%200.18#0>,
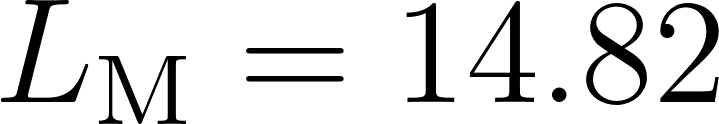
<https://www.codecogs.com/eqnedit.php?latex=L_%5Ctext%7BM%7D%20%3D%2014.82#0> Morgans (*27*,*119*), and
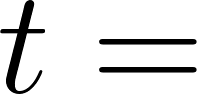
<https://www.codecogs.com/eqnedit.php?latex=t%20%3D%20#0> 6,200 generations, the standard deviation of
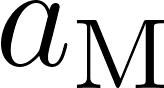
<https://www.codecogs.com/eqnedit.php?latex=a_%5Ctext%7BM%7D#0> is
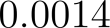
<https://www.codecogs.com/eqnedit.php?latex=0.0014#0>.

To convert this to a variance of the introgressed fraction of the physical (bp) genome, note first that the distribution of block number does not depend on whether block length is measured in Morgans or bp. Let
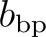
<https://www.codecogs.com/eqnedit.php?latex=b_%5Ctext%7Bbp%7D#0> and
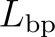
<https://www.codecogs.com/eqnedit.php?latex=L_%5Ctext%7Bbp%7D#0> be the bp lengths of an introgressed block and of the haploid genome respectively, and
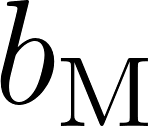
<https://www.codecogs.com/eqnedit.php?latex=b_%5Ctext%7BM%7D#0> and
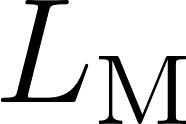
<https://www.codecogs.com/eqnedit.php?latex=L_%5Ctext%7BM%7D#0> their Morgan lengths. The relationship between the bp and Morgan length of a block is
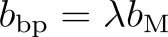
<https://www.codecogs.com/eqnedit.php?latex=b_%5Ctext%7Bbp%7D%20%3D%20%5Clambda%20b_%5Ctext%7BM%7D#0>, where
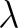
<https://www.codecogs.com/eqnedit.php?latex=%5Clambda#0> (with units bp/Morgan) varies across the genome and is independent of
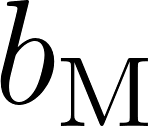
<https://www.codecogs.com/eqnedit.php?latex=b_%5Ctext%7BM%7D#0>.

We have

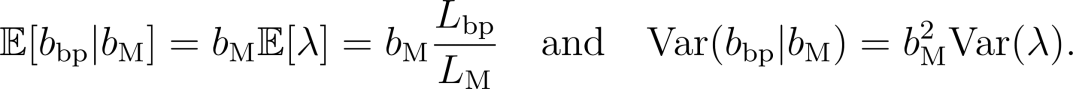
<https://www.codecogs.com/eqnedit.php?latex=%5Cmathbb%7BE%7D%5Bb_%5Ctext%7Bbp%7D%7Cb_%5Ctext%7BM%7D%5D%20%3D%20b_%5Ctext%7BM%7D%5Cmathbb%7BE%7D%5B%5Clambda%5D%20%3D%20b_%5Ctext%7BM%7D%20%5Cfrac%7BL_%5Ctext%7Bbp%7D%7D%7BL_%5Ctext%7BM%7D%7D%20%5C%2C%5C%2C%5C%2C%5C%2C%20%5Ctext%7B%20and%20%7D%20%5C%2C%5C%2C%5C%2C%5C%2C%20%5Ctext%7BVar%7D(b_%5Ctext%7Bbp%7D%20%7C%20b_%5Ctext%7BM%7D)%20%3D%20b_%5Ctext%7BM%7D%5E2%20%5Ctext%7BVar%7D(%5Clambda).#0>

From the law of iterated expectations,

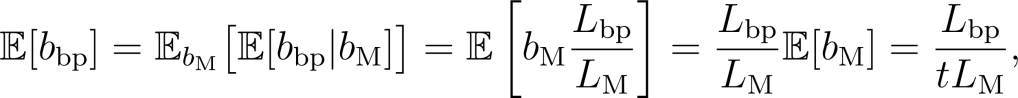
<https://www.codecogs.com/eqnedit.php?latex=%5Cmathbb%7BE%7D%5Bb_%5Ctext%7Bbp%7D%5D%20%3D%20%5Cmathbb%7BE%7D_%7Bb_%5Ctext%7BM%7D%7D%5Cbig%5B%20%5Cmathbb%7BE%7D%5Bb_%5Ctext%7Bbp%7D%7Cb_%5Ctext%7BM%7D%5D%20%5Cbig%5D%20%3D%20%5Cmathbb%7BE%7D%5Cleft%5B%20b_%5Ctext%7BM%7D%20%5Cfrac%7BL_%5Ctext%7Bbp%7D%7D%7BL_%5Ctext%7BM%7D%7D%20%5Cright%5D%20%3D%20%5Cfrac%7BL_%5Ctext%7Bbp%7D%7D%7BL_%5Ctext%7BM%7D%7D%5Cmathbb%7BE%7D%5Bb_%5Ctext%7BM%7D%5D%20%3D%20%5Cfrac%7BL_%5Ctext%7Bbp%7D%7D%7BtL_%5Ctext%7BM%7D%7D%2C#0>

and from the law of total variance,

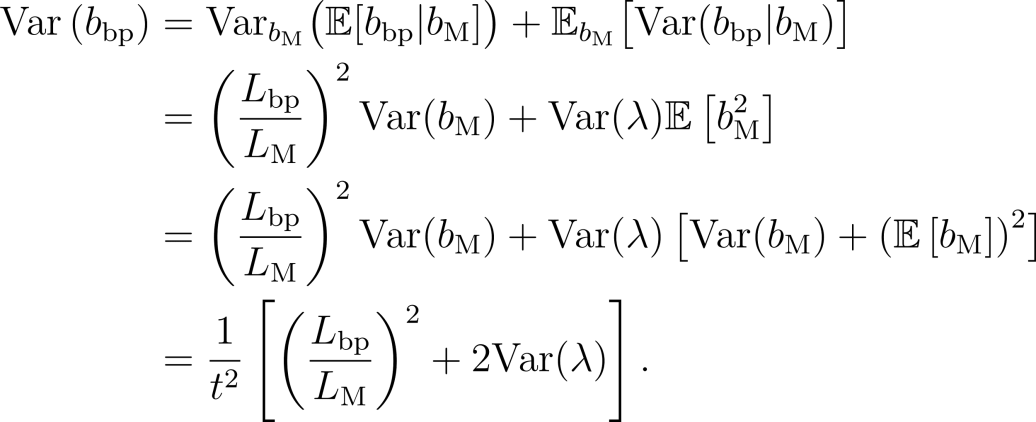
<https://www.codecogs.com/eqnedit.php?latex=%5Cbegin%7Balign*%7D%5Ctext%7BVar%7D%20%5Cleft(%20b_%5Ctext%7Bbp%7D%20%5Cright)%20%26%3D%20%5Ctext%7BVar%7D_%7Bb_%5Ctext%7BM%7D%7D%5Cbig(%20%20%5Cmathbb%7BE%7D%5Bb_%5Ctext%7Bbp%7D%7Cb_%5Ctext%7BM%7D%5D%20%5Cbig)%20%2B%20%20%5Cmathbb%7BE%7D_%7Bb_%5Ctext%7BM%7D%7D%5Cbig%5B%20%20%20%5Ctext%7BVar%7D(b_%5Ctext%7Bbp%7D%20%7C%20b_%5Ctext%7BM%7D)%20%20%20%5Cbig%5D%5C%5C%5C%5C%20%26%3D%20%5Cleft(%5Cfrac%7BL_%5Ctext%7Bbp%7D%7D%7BL_%5Ctext%7BM%7D%7D%5Cright)%5E2%5Ctext%7BVar%7D(b_%5Ctext%7BM%7D)%20%2B%20%5Ctext%7BVar%7D(%5Clambda)%5Cmathbb%7BE%7D%5Cleft%5B%20%20b_%5Ctext%7BM%7D%5E2%20%20%5Cright%5D%5C%5C%5C%5C%20%26%3D%20%5Cleft(%5Cfrac%7BL_%5Ctext%7Bbp%7D%7D%7BL_%5Ctext%7BM%7D%7D%5Cright)%5E2%5Ctext%7BVar%7D(b_%5Ctext%7BM%7D)%20%2B%20%20%5Ctext%7BVar%7D(%5Clambda)%5Cleft%5B%20%5Ctext%7BVar%7D(b_%5Ctext%7BM%7D)%20%2B%20%5Cleft(%5Cmathbb%7BE%7D%5Cleft%5Bb_%5Ctext%7BM%7D%5Cright%5D%5Cright)%5E2%20%5Cright%5D%5C%5C%5C%5C%20%26%3D%20%5Cfrac%7B1%7D%7Bt%5E2%7D%5Cleft%5B%20%5Cleft(%5Cfrac%7BL_%5Ctext%7Bbp%7D%7D%7BL_%5Ctext%7BM%7D%7D%5Cright)%5E2%20%2B%202%5Ctext%7BVar%7D(%5Clambda)%5Cright%5D.%5Cend%7Balign*%7D#0>

The appropriate scale at which
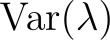
<https://www.codecogs.com/eqnedit.php?latex=%5Ctext%7BVar%7D(%5Clambda)#0> should be calculated is
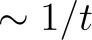
<https://www.codecogs.com/eqnedit.php?latex=%5Csim%201%2Ft#0> Morgans.

Conditional on receiving
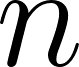
<https://www.codecogs.com/eqnedit.php?latex=n#0> blocks, an individual's introgressed fraction
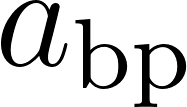
<https://www.codecogs.com/eqnedit.php?latex=a_%5Ctext%7Bbp%7D#0> obeys

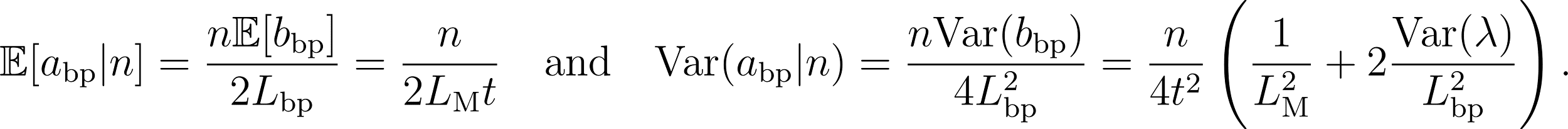
<https://www.codecogs.com/eqnedit.php?latex=%5Cmathbb%7BE%7D%5Ba_%5Ctext%7Bbp%7D%7Cn%5D%20%3D%20%5Cfrac%7Bn%5Cmathbb%7BE%7D%5Bb_%5Ctext%7Bbp%7D%5D%7D%7B2L_%5Ctext%7Bbp%7D%7D%20%3D%20%5Cfrac%7Bn%7D%7B2L_%5Ctext%7BM%7Dt%7D%20%5C%2C%5C%2C%5C%2C%5C%2C%20%5Ctext%7B%20and%20%7D%20%5C%2C%5C%2C%5C%2C%5C%2C%20%5Ctext%7BVar%7D(a_%5Ctext%7Bbp%7D%7Cn)%20%3D%20%5Cfrac%7Bn%5Ctext%7BVar%7D(b_%5Ctext%7Bbp%7D)%7D%7B4L_%5Ctext%7Bbp%7D%5E2%7D%20%3D%20%5Cfrac%7Bn%7D%7B4t%5E2%7D%5Cleft(%20%20%5Cfrac%7B1%7D%7BL_%5Ctext%7BM%7D%5E2%7D%20%2B%202%5Cfrac%7B%5Ctext%7BVar%7D(%5Clambda)%7D%7BL_%5Ctext%7Bbp%7D%5E2%7D%20%5Cright).#0>

From the law of total variance,

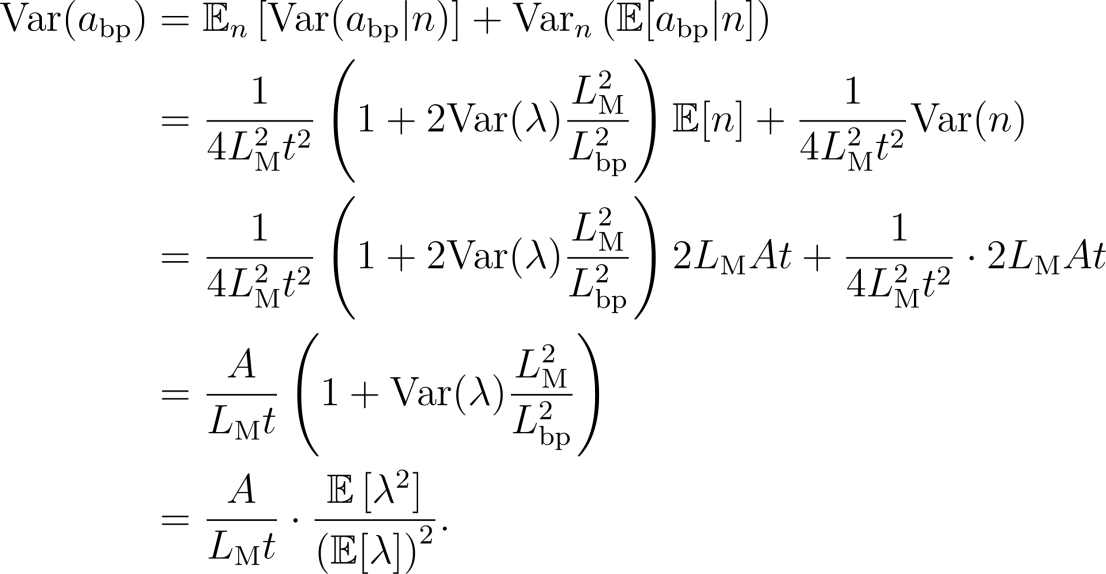
<https://www.codecogs.com/eqnedit.php?latex=%5Cbegin%7Balign*%7D%5Ctext%7BVar%7D(a_%5Ctext%7Bbp%7D)%20%26%3D%20%5Cmathbb%7BE%7D_n%5Cleft%5B%20%5Ctext%7BVar%7D(a_%5Ctext%7Bbp%7D%7Cn)%20%5Cright%5D%20%2B%20%5Ctext%7BVar%7D_n%5Cleft(%5Cmathbb%7BE%7D%5Ba_%5Ctext%7Bbp%7D%7Cn%5D%5Cright)%5C%5C%5C%5C%20%26%3D%5Cfrac%7B1%7D%7B4L_%5Ctext%7BM%7D%5E2t%5E2%7D%5Cleft(%201%20%2B%202%5Ctext%7BVar%7D(%5Clambda)%5Cfrac%7BL_%5Ctext%7BM%7D%5E2%7D%7BL_%5Ctext%7Bbp%7D%5E2%7D%20%5Cright)%5Cmathbb%7BE%7D%5Bn%5D%20%2B%20%5Cfrac%7B1%7D%7B4L_%5Ctext%7BM%7D%5E2t%5E2%7D%5Ctext%7BVar%7D(n)%5C%5C%5C%5C%20%26%3D%5Cfrac%7B1%7D%7B4L_%5Ctext%7BM%7D%5E2t%5E2%7D%5Cleft(%201%20%2B%202%5Ctext%7BVar%7D(%5Clambda)%5Cfrac%7BL_%5Ctext%7BM%7D%5E2%7D%7BL_%5Ctext%7Bbp%7D%5E2%7D%20%5Cright)%202L_%5Ctext%7BM%7DAt%20%2B%20%5Cfrac%7B1%7D%7B4L_%5Ctext%7BM%7D%5E2t%5E2%7D%5Ccdot%202L_%5Ctext%7BM%7DAt%5C%5C%5C%5C%20%26%3D%20%20%5Cfrac%7BA%7D%7BL_%5Ctext%7BM%7Dt%7D%5Cleft(%20%201%20%2B%20%20%20%20%5Ctext%7BVar%7D(%5Clambda)%5Cfrac%7BL_%5Ctext%7BM%7D%5E2%7D%7BL_%5Ctext%7Bbp%7D%5E2%7D%20%20%20%20%5Cright)%5C%5C%5C%5C%20%26%3D%20%20%5Cfrac%7BA%7D%7BL_%5Ctext%7BM%7Dt%7D%5Ccdot%20%5Cfrac%7B%5Cmathbb%7BE%7D%5Cleft%5B%5Clambda%5E2%5Cright%5D%7D%7B%5Cleft(%5Cmathbb%7BE%7D%5B%5Clambda%5D%5Cright)%5E2%7D.%5Cend%7Balign*%7D#0>

So the variance across individuals in the fraction of their physical genome that is introgressed is larger than the analogous calculation in Morgans by a factor of
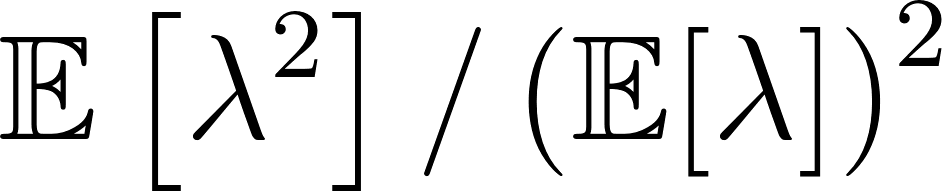
<https://www.codecogs.com/eqnedit.php?latex=%5Cmathbb%7BE%7D%5Cleft%5B%5Clambda%5E2%5Cright%5D%2F%5Cleft(%5Cmathbb%7BE%7D%5B%5Clambda%5D%5Cright)%5E2#0>. Since
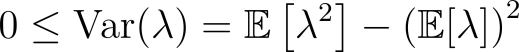
<https://www.codecogs.com/eqnedit.php?latex=0%20%5Cleq%20%5Ctext%7BVar%7D(%5Clambda)%20%3D%20%5Cmathbb%7BE%7D%5Cleft%5B%5Clambda%5E2%5Cright%5D%20-%20%5Cleft(%5Cmathbb%7BE%7D%5B%5Clambda%5D%5Cright)%5E2#0>, this factor is greater than or equal to 1, being equal to 1 only if
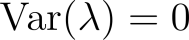
<https://www.codecogs.com/eqnedit.php?latex=%5Ctext%7BVar%7D(%5Clambda)%3D0#0>, i.e., if the recombination rate is uniform across the genome.

To calculate
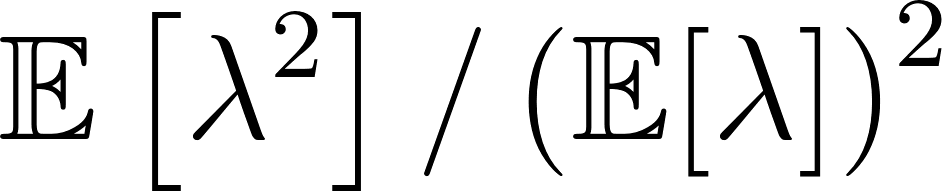
<https://www.codecogs.com/eqnedit.php?latex=%5Cmathbb%7BE%7D%5Cleft%5B%5Clambda%5E2%5Cright%5D%2F%5Cleft(%5Cmathbb%7BE%7D%5B%5Clambda%5D%5Cright)%5E2#0>, we use a linkage map ported to the B73 v4 reference genome (*27*,*119*). Dividing the maize genome into segments of length
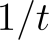
<https://www.codecogs.com/eqnedit.php?latex=1%2Ft#0> Morgans, with
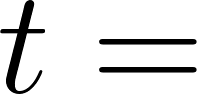
<https://www.codecogs.com/eqnedit.php?latex=t%3D#0> 6,200, and calculating
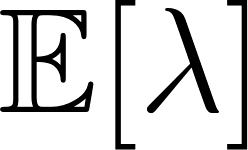
<https://www.codecogs.com/eqnedit.php?latex=%5Cmathbb%7BE%7D%5B%5Clambda%5D#0> and
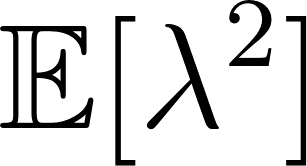
<https://www.codecogs.com/eqnedit.php?latex=%5Cmathbb%7BE%7D%5B%5Clambda%5E2%5D#0> at this scale, we find that

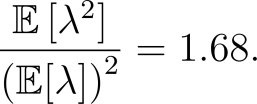
<https://www.codecogs.com/eqnedit.php?latex=%5Cfrac%7B%5Cmathbb%7BE%7D%5Cleft%5B%5Clambda%5E2%5Cright%5D%7D%7B%5Cleft(%5Cmathbb%7BE%7D%5B%5Clambda%5D%5Cright)%5E2%7D%20%3D%201.68.#0>

(Reassuringly, this value is not particularly sensitive to changes in the Morgan scale at which
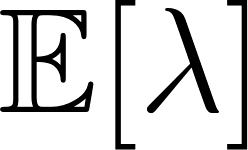
<https://www.codecogs.com/eqnedit.php?latex=%5Cmathbb%7BE%7D%5B%5Clambda%5D#0> and
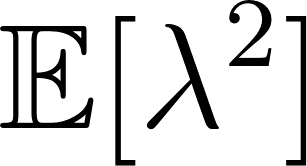
<https://www.codecogs.com/eqnedit.php?latex=%5Cmathbb%7BE%7D%5B%5Clambda%5E2%5D#0> are calculated. For example, if the scale is ten-fold larger (
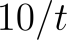
<https://www.codecogs.com/eqnedit.php?latex=10%2Ft#0>), the variance inflation factor is
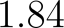
<https://www.codecogs.com/eqnedit.php?latex=1.84#0>, while if the scale is ten-fold smaller (
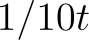
<https://www.codecogs.com/eqnedit.php?latex=1%2F10t#0>), the variance inflation factor is
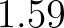
<https://www.codecogs.com/eqnedit.php?latex=1.59#0>.)

So the standard deviation of
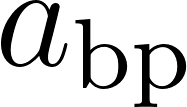
<https://www.codecogs.com/eqnedit.php?latex=a_%5Ctext%7Bbp%7D#0> is about
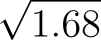
<https://www.codecogs.com/eqnedit.php?latex=%5Csqrt%7B1.68%7D#0> larger than the standard deviation of
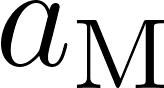
<https://www.codecogs.com/eqnedit.php?latex=a_%5Ctext%7BM%7D#0>, owing to heterogeneity in the recombination rate across the genome. If
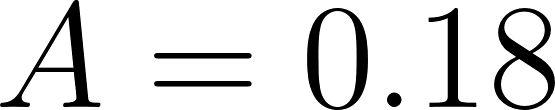
<https://www.codecogs.com/eqnedit.php?latex=A%20%3D%200.18#0>,
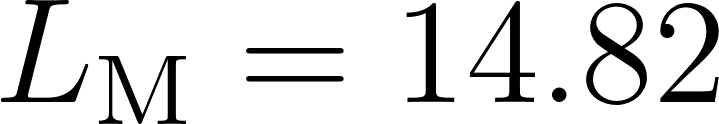
<https://www.codecogs.com/eqnedit.php?latex=L_%5Ctext%7BM%7D%20%3D%2014.82#0> Morgans,
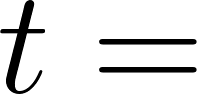
<https://www.codecogs.com/eqnedit.php?latex=t%20%3D%20#0> 6,200 generations, and
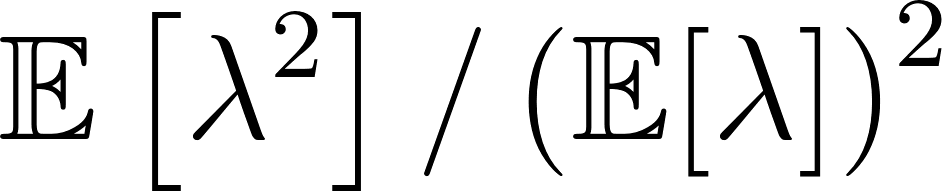
<https://www.codecogs.com/eqnedit.php?latex=%5Cmathbb%7BE%7D%5Cleft%5B%5Clambda%5E2%5Cright%5D%2F%5Cleft(%5Cmathbb%7BE%7D%5B%5Clambda%5D%5Cright)%5E2#0>= 1.68, we calculate that, under a model with no additional admixture or selection, the standard deviation across individuals of the fraction of their genomes that are introgressed,
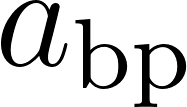
<https://www.codecogs.com/eqnedit.php?latex=a_%5Ctext%7Bbp%7D#0> should be about
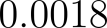
<https://www.codecogs.com/eqnedit.php?latex=0.0018#0>.

Introgression at *Inv1n*

We observed an obvious elevation of *mexicana* dosage in the *Inv1n* (*63*) region. One explanation is the homologous region of *Inv1n* in maize was introgressed from *mexicana*. Another possible explanation, however, is that *mexicana*, maize and *parviglumis* without *Inv1n* have the same ancestral variant while the *Inv1n* had increased in frequency in *parviglumis* due to selection. If the later explanation was correct, admixture analysis using only *parviglumis* without *Inv1n* should reveal low *mexicana* ancestry in maize. Using inversion genotypes at 8 regions within the inversion from Chen *et al.* (*39*), we reran our ELAI analysis on chromosome 1 with *parviglumis* individuals homozygous for the uninverted allele. In this analysis we still found a clear elevation of *mexicana* ancestry compared to the rest of the genome (**fig. S19**).

Overexpression and knovkout of *ZmPRR37a* in maize

The 1,689 bp full-length coding sequence (CDS) of ZmPRR37a_T03 (annotation version:B73 5b+) was amplified from the cDNA library of maize inbred line B73. In order to distinguish between endogenous gene and transgenic sequences, the CDS of ZmPRR37a_T03 was codon-optimized, infused with 3×FLAG tag on the N terminus and then synthesized by GenScript Inc. (NanJing, China). The CoZmPRR37a_T03 (codon-optimized ZmPRR37a_T03) was inserted into the modified binary vector pCAMBIA3300 driven by the ZmUbi promoter for the overexpression study (*120*) . These binary vector harboring CoZmPRR37a_T03 were transformed into maize inbred line KN5585 by Agrobacterium tumefaciens-mediated method as described (*67*) . All of the transgenic plants were identified by herbicide tests, PCR validations of bar and CoZmPRR37a_T03. (The primers for bar and CoZmPRR37a_T03 are shown in **table S11**). The T0 transgenic plants (CoZmPRR37a_T03-OE) were backcrossed to receptor KN5585 to produce the transgene-positive and transgene-negative segregants in T1, T2 and T3 generations.

### For qRT-PCR analysis, leaf tissues were collected at stage V6, and RNA was isolated from each tissue with EasyPure Plant RNA Kit (TransGen, Beijing, China). The cDNA was synthesized using a TransScript One-Step gDNA Removal and cDNA Synthesis SuperMix (TransGen, Beijing, China). The expression of CoZmPRR37a_T03 was determined by real-time PCR using primers qCoZmPRR37a_T03 (FastStart Essential DNA Green Master, Roche, Mannheim, Germany), and ACTIN primers as an endogenous control. Non-transgenic plants were used as negative control. Quantitative real-time PCR using SYBR Green PCR mix (FastStart Essential DNA Green Master, Roche, Mannheim, Germany) was performed with an ABI QuantStudio 3 system according to the manufacturer's’ instructions. Each sample is set with 3 technical repetitions. It is worth noting that although the same protein can be expressed, the nucleotide sequences of CoZmPRR37a_T03 and ZmPRR37a_T03 are significantly different due to codon-optimization. So the primers of qRT-PCR can not recognize native ZmPRR37a_T03 genes.

The transgene-positive and transgene-negative segregants of CoZmPRR37a_T03-OE (T2 and T3 generation) were evaluated for flowering time and other agronomic traits in short-day (Winter of 2019 in Foluo town, Ledong City, Hainan Province (18°34′N 108°43′E)) and long-day (Summer of 2022 in Gongzhuling city, Jilin province (43°30′N 124°49′E)) environments. The flowering time traits of each individual, eg. DTT: Days to Tassel (days), DTA: Days to Anthesis (days), DTS: Days to Silking (days) were measured.

Knockout mutants of *ZmPRR37a* were produced from a high-throughput genome-editing system (*67*). A double sgRNAs pool approach was used for vector construction and KN5585 was transformed (Target1: 5’-GCAGGCAGAAATAGTCATCCTGG-3’; Target2: 5’- GAGTACAGCTGGTGCGAGCACGG-3’). The genotype of lines was identified by PCR amplification and sanger sequencing by target-specific primers (F: 5’-TAACGAGAGGGTGATGTTGCC-3’; R: 5’-AGCTGTTCTTGTCGTGGTGTC-3’). The homozygous gene-edited lines of T1 generation and the receptor KN5585 were investigated for flowering time, which including days to tasseling (DTT), days to anthesis (DTA), and days to silking (DTS), at Gongzhuling city, Jilin province (N 43° 30′, E 124° 49′) in 2018. And the segregating populations were constructed by crossing homozygous gene-edited lines with KN5585 then do selfing. The homozygous gene-edited lines and wild-type lines from segregating population were investigated for flowering time at Foluo town, Ledong City, Hainan Province (N 18° 34′, E 108° 43′) in 2020.

Historical morphological admixture

Varying levels of teosinte introgression in the morphology of the maize ear has been used as a character to distinguish landraces (*69*). Wellhausen *et al.*(*69*) characterize teosinte introgression using a categorical scale from 0 to 4, largely summarizing rachis induration and the length and number of hairs on the glume. To associate historical measurements of landraces to our contemporary STRUCTURE teosinte ancestry values, we collected the `primary_landrace` identifier from CIMMYT GRIN, and annotated our landrace samples with the phenotypic values listed in Table 16 of Wellhausen 1952 (**fig. S23**). The correlation between *parviglumis* admixture and historical measurements of introgression in these 1698 maize samples is positive, statistically significant, and large (**table S6**, Spearman’s ρ=0.67, S = 2.67e+08, p<0.001).

Admixture Mapping in the Inbred Diversity Panel

We downloaded 614 phenotypes that have been measured on a large maize inbred association panel (*43*,*90*), including agronomic, disease, metabolomic, and kernel biochemistry traits. These were filtered to remove traits with >10% missing data or simple binary phenotypes, subsampled down to a set of 33 phenotypes for which all pairwise correlations were less than 0.6 (**fig. S29**), and log10-transformed. We then performed multivariate GWAS using the JointGWAS R package. JointGWAS tests each marker for an association between admixture proportion and *any* of the 33 traits using a generalized least squares F-test with a pre-estimated variance-covariance matrix accounting for genetic and non-genetic covariances among traits (due to pleiotropy and environmental covariation) and among genotypes (due to shared alleles). This covariance matrix was first estimated using the R package MegaLMM (*121*) using a model with a single random effect with covariance proportional to the whole-genome kinship matrix among genotypes, estimated from centered identity-by-state comparisons using 1 million random SNPs in Tassel 5 (*122*). We specified 50 latent factors with the Automatic Relevance Determination prior for the factor loadings with degree of freedom parameter set to 2, and an expected decline of 50% importance between consecutive factors (δ_2_ ~ Ga(3,2)), and a prior that assigned 50% probability to the heritability of each factor being 0.98 and 50% probability that the heritability < 0.98, reflecting the fact that most traits were measured on different individuals so non-genetic covariances should be rare. We extracted the posterior means of the genetic and non-genetic covariances from a single model run in which 1000 posterior samples were collected at a thinning rate of 2 after a burnin of 2000 iterations. Given these estimates we formed an estimate of the full covariance of the data using the JointGWAS (*80*) function make_cholL_Sigma_inv, and ran the F-test on all loci with at least 5% inferred mexicana alleles using the JointGWAS function EMMAX_ANOVA, allowing an intercept for each of the 33 traits and a trait-specific marker effect. The F-test tests the null hypothesis that the marker effect equals zero for all traits.

Admixture mapping across environments in traditional varieties

We used trait data from the F1 progeny of approximately 4,500 traditionally cultivated varieties crossed to testers crossed to testers and grown in 13 trials (*81*). Within each trial, trait data were reported as BLUPs accounting for spatial variation across the field. To facilitate comparisons across fields we de-regressed the BLUPs within each trial by dividing each value by the average reliability (1-average BLUP variance/genetic variance). We eliminated BLUPs from testers crossed to fewer than 20 accessions per trial. Then for each trait we ran a GWAS testing for an association between the admixture proportion at each marker and each trait in any trial again using the JointGWAS R package . For each trait we estimated the genetic and non-genetic covariances among trials using MegaLMM as described above, except the number of factors was set to one fewer than the number of trials. We again used the make_cholL_Sigma_inv to calculate the full covariance matrix among observations and tested each marker using the generalized least squares F-test using the EMMAX_ANOVA function for all markers with at least 1% frequency of the *mexicana* allele.

Variance partitioning

We partitioned variance in phenotypes between two kinship matrices. The first was the IBS kinship matrix from SNP data used in our GWAS. We then estimated a second kinship matrix based on ELAI admixture scores using OSCA (*123*), with the commands ‘osca_Linux --efile amp.elai.txt --methylation-beta --make-bod --no-fid -lower-beta 0.001 --out elai.kinship’ and ‘osca_Linux --befile elai.kinship --make-orm-gz --out elai.orm’. We used LDAK (*124*) to calculate the variance explained by each kinship matrix, imputing the missing data and constraining heritabilities to be between 0 and 1. The command used was ‘ldak5.2.linux --reml --reml ./results/standard/standard --mgrm ./kinship_standard.txt --pheno ./phenos2_log_ldak.txt --mpheno -1 –constrain YES –dentist YES --kinship-details NO’. For each phenotype we ran the model with or without the “he-starts” flag (which uses Haseman-Elston to estimate a starting value for the REML) and recorded results for the model with the highest restricted likelihood. To test for the accuracy of this approach, we used these kinship matrices to simulate traits covering a grid of 12 combinations of heritabilities distributed between the two kinship matrices and independent error. We drew three random vectors with covariances proportional to the *mexicana* introgression kinship matrix, the background kinship matrix, and the identity matrix, respectively, and then re-weighted each by the assigned variance component proportions. We then summed the three re-weighted vectors to form a simulated trait with appropriate variance and covariance. The total heritability varied at 0.2, 0.5, or 0.8 and, under each, the *mexicana* introgression made up 10%, 25%, 50%, or 75% of the total heritability. For the code used to simulate these traits, see https://github.com/rossibarra/maize_origins/simulate_pheontypes.Rmd. From there the heritability for the rest of the genome and error were calculated. These simulated trait values were run through the same REML solver in LDAK as above (*124*), constraining values to be between 0 and 1 and using Haseman-Elston regression to choose starting values. While there was considerable variability for simulated traits with lower heritability, across all parameter values the means of the 10 replicates were very close to the true values (**fig. S30**).

**Supplementary Tables**

Table S1. Basic information regarding the sampled accessions.

Table S2. Sequencing and mapping profile of 338 samples newly sequenced in this study.

Table S3. *f4* statistic of all samples in Fig 1.

Table S4. *mexicana* dosage estimated by ELAI.

Table S5. The fixed *mexicana* alleles in maize population.

Table S6. *parviglumis* admixture and historical measurements (*69*) of introgression

Table S7. The log transformed phenotypes used for GWAS.

Table S8. Candidate genes of multivariate admixture mapping in the maize inbred association panel.

Table S9. The proportion of additive genetic variance (narrow sense heritability) contributed by *mexicana* across a set of 33 phenotypes.

Table S10. The reference panel for ELAI analysis.

Table S11. Primers used in this study.

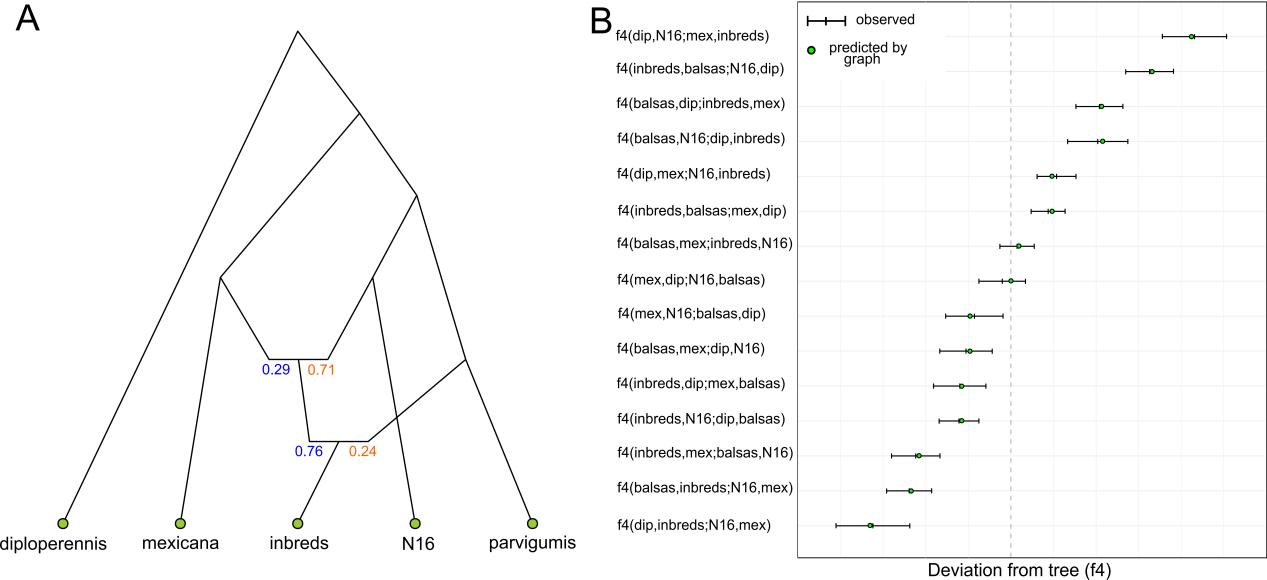

**Fig. S1.** (**A**) Admixture graph of modern maize inbreds (inbreds) and teosinte. Colors in blue and yellow at each node represent the estimated proportion of admixture from each source population. “inbreds” refers to 507 modern inbred lines (**table S1**) (*43*).(**B**) Fit of observed f4 statistics with the graph. Observed estimates of f4 are shown as short vertical lines with error bars (black lines) while circles show the expected values generated by the graph that are within (green) or outside of (red) the range observed in the data.

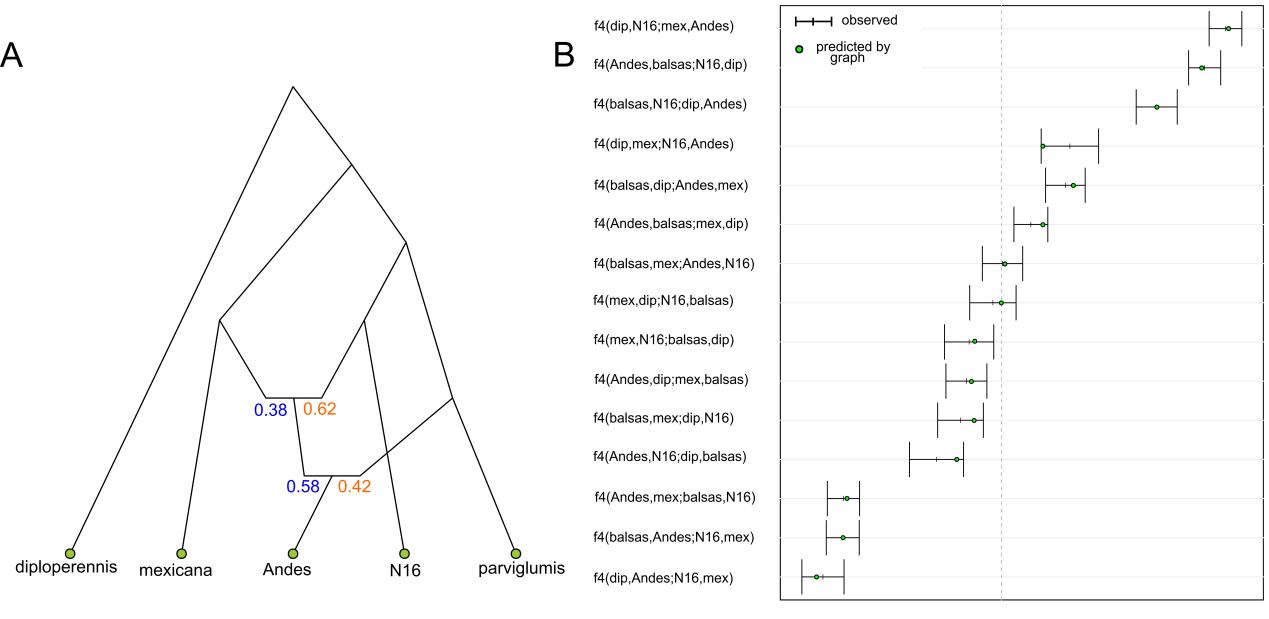

**Fig. S2.** (**A**) Admixture graph of landraces (Andes) and teosinte. Colors in blue and yellow at each node represent the estimated proportion of admixture from each source population. “Andes” refers to the Peruvian Andes landrace population including 5 individuals **(table S1**) (*40*). (**B**) Fit of observed f4 statistics with the graph. Observed estimates of f4 are shown as short vertical lines with error bars (black lines) while circles show the expected values generated by the graph that are within (green) or outside of (red) the range observed in the data.

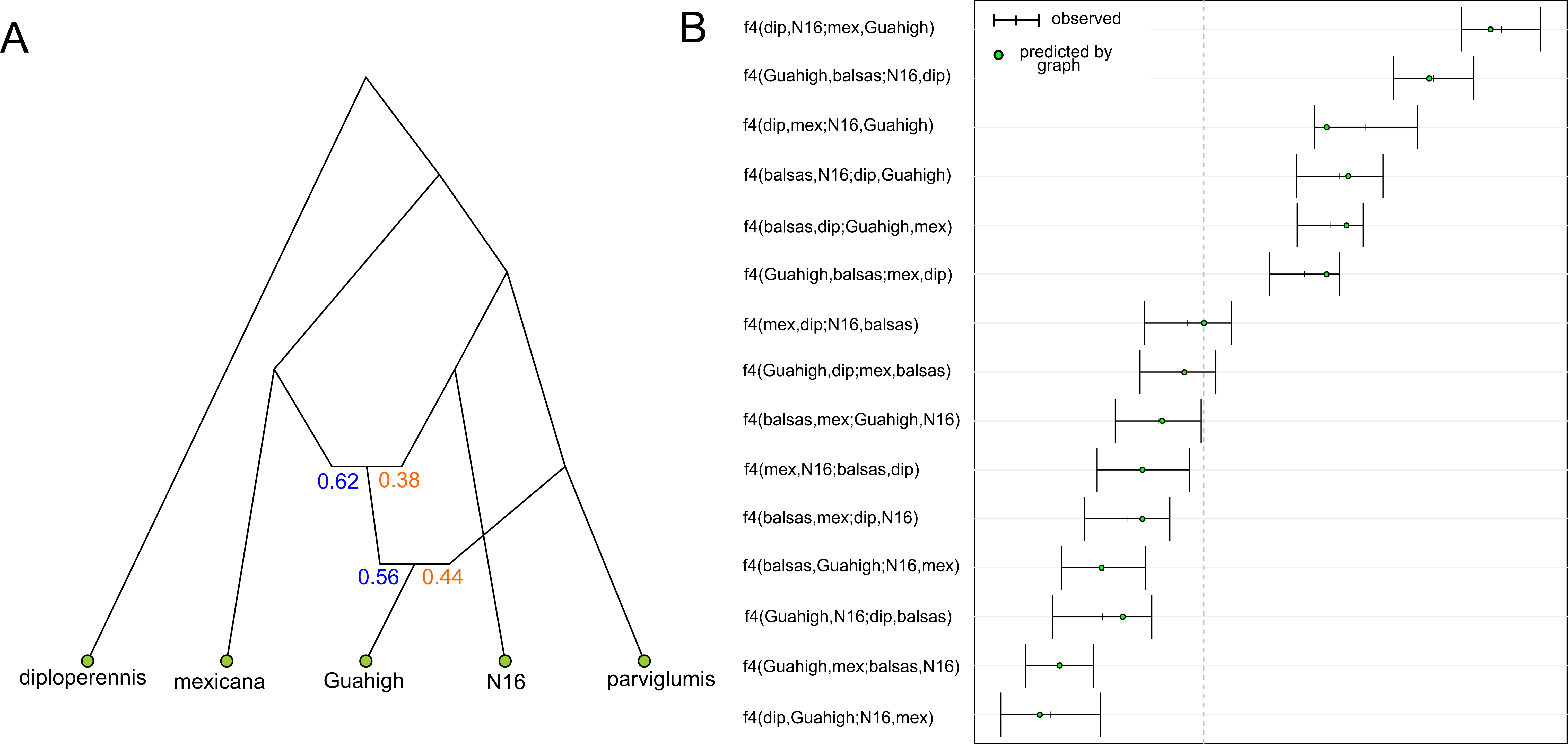

**Figure S3** (**A**) Admixture graph of landraces (Guahigh) and teosinte. Colors in blue and yellow at each node represent the estimated proportion of admixture from each source population. “Guahigh” refers to the Guatemalan highlands landrace population including 3 individuals (**table S1**) (*40*). (**B**) Fit of observed f4 statistics with the graph. Observed estimates of f4 are shown as short vertical lines with error bars (black lines) while circles show the expected values generated by the graph that are within (green) or outside of (red) the range observed in the data.

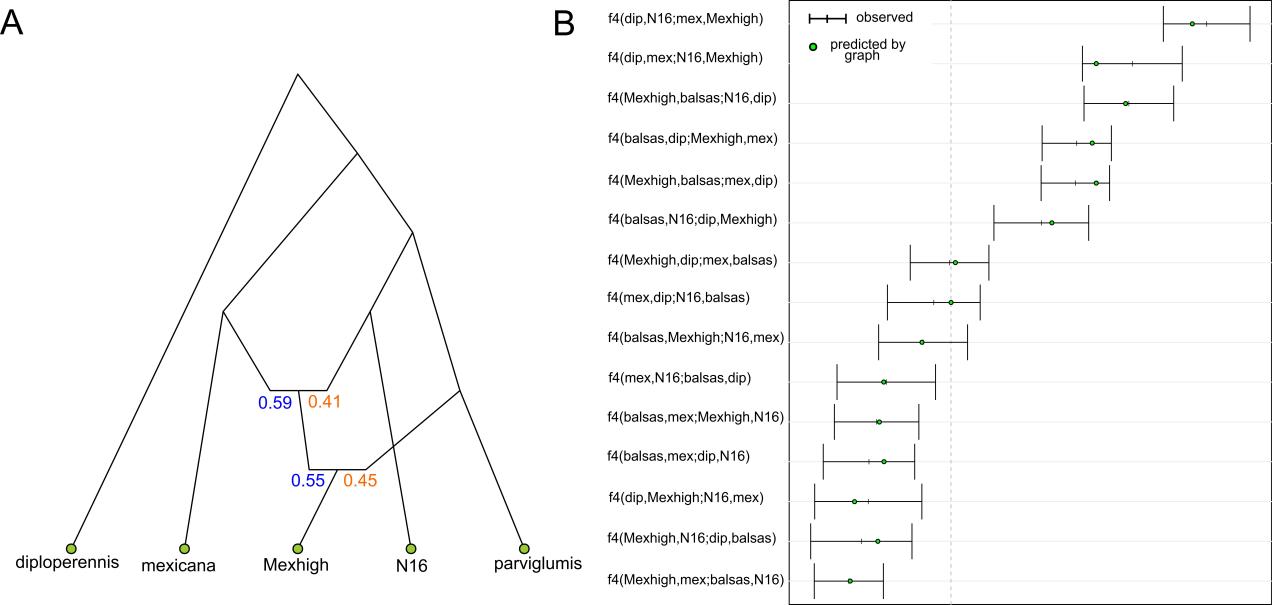

**Figure S4** (**A**) Admixture graph of landraces (Mexhigh) and teosinte. Colors in blue and yellow at each node represent the estimated proportion of admixture from each source population. “Mexhigh” refers to the Mexican highlands landrace population including 5 individuals (**table S1**) (*40*). (**B**) Fit of observed f4 statistics with the graph. Observed estimates of f4 are shown as short vertical lines with error bars (black lines) while circles show the expected values generated by the graph that are within (green) or outside of (red) the range observed in the data.

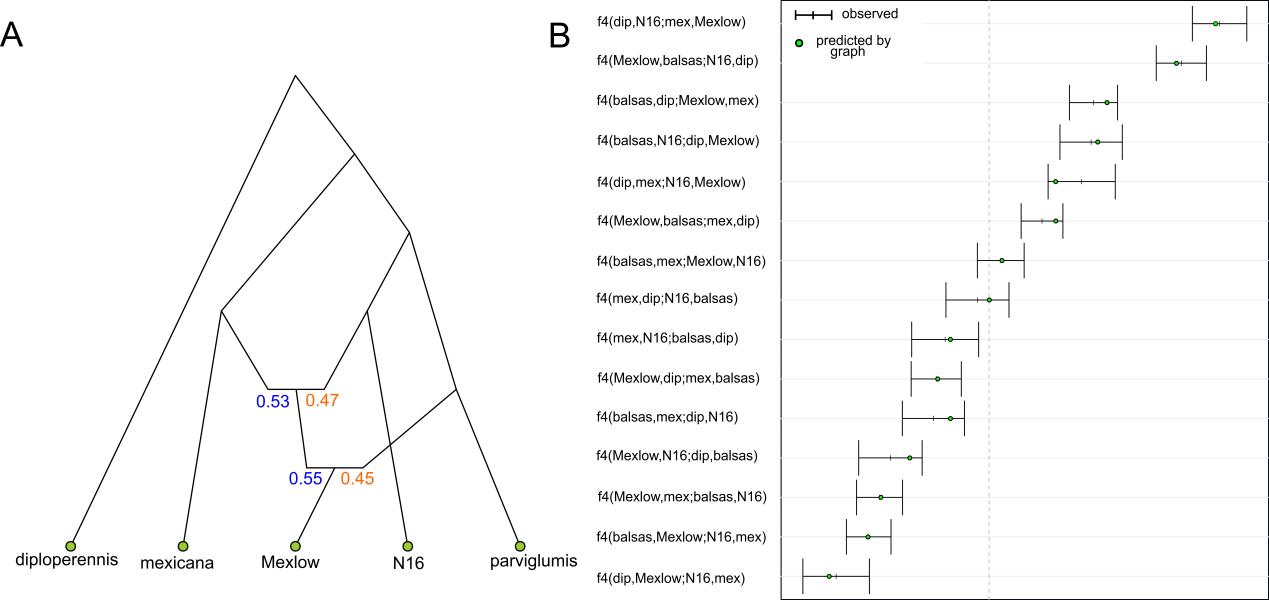

**Figure S5** (**A**) Admixture graph of landraces (Mexlow) and teosinte. Colors in blue and yellow at each node represent the estimated proportion of admixture from each source population. “Mexlow” refers to the Mexican lowlands landrace population including 5 individuals (**table S1**) (*40*). (**B**) Fit of observed f4 statistics with the graph. Observed estimates of f4 are shown as short vertical lines with error bars (black lines) while circles show the expected values generated by the graph that are within (green) or outside of (red) the range observed in the data.

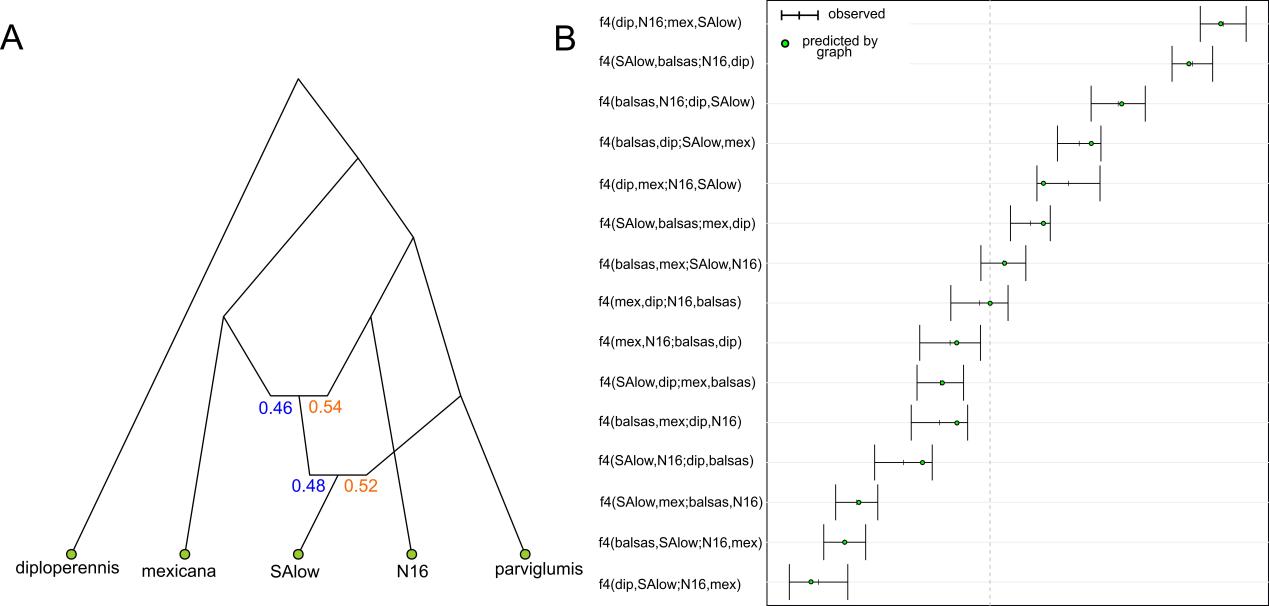

**Fig. S6.** (**A**) Admixture graph of landraces (SA_lowlands) and teosinte. Colors in blue and yellow at each node represent the estimated proportion of admixture from each source population. “SA_lowlands” refers to the South American lowlands landrace population including 6 individuals (**table S1**) (*40*). (**B**) Fit of observed f4 statistics with the graph. “Balsas” refers to the Balsas samples of *parviglumis.* Observed estimates of f4 are shown as short vertical lines with error bars (black lines) while circles show the expected values generated by the graph that are within (green) or outside of (red) the range observed in the data.

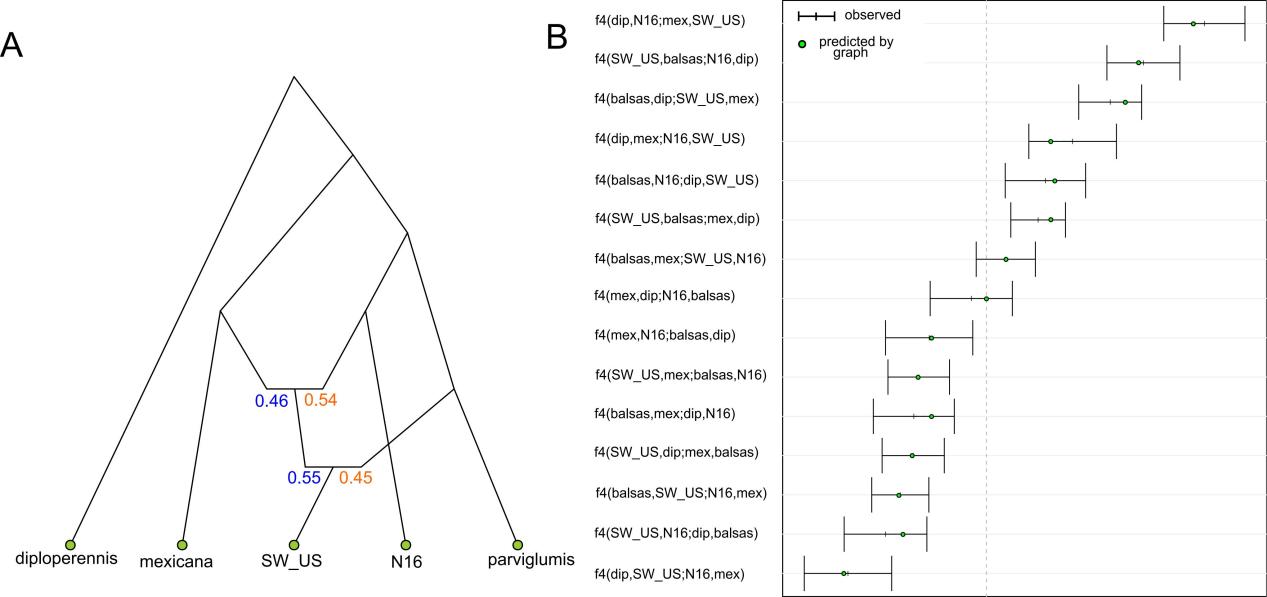

**Figure S7** (**A**) Admixture graph of landraces (SW_US) and teosinte. Colors in blue and yellow at each node represent the estimated proportion of admixture from each source population. “SW_US” refers to the southwestern US highlands landrace population including 6 individuals (**table S1**) (*40*). (**B**) Fit of observed f4 statistics with the graph. Observed estimates of f4 are shown as short vertical lines with error bars (black lines) while circles show the expected values generated by the graph that are within (green) or outside of (red) the range observed in the data.

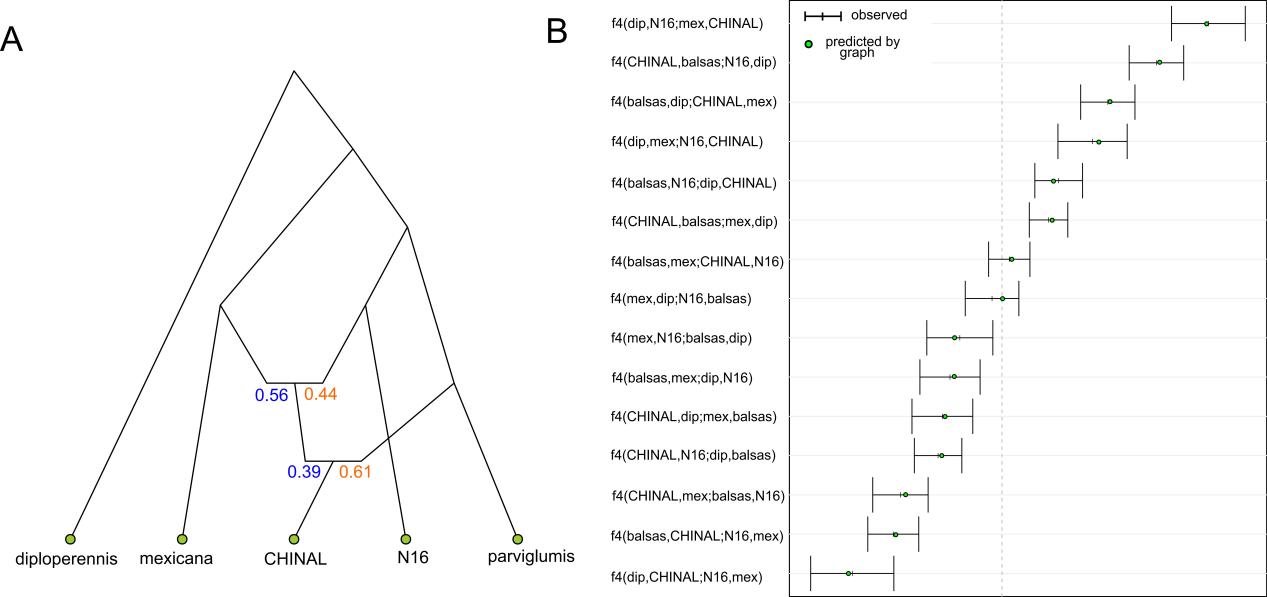

**Figure S8** (**A**) Admixture graph of landraces (CHINAL) and teosinte. Colors in blue and yellow at each node represent the estimated proportion of admixture from each source population. “CHINAL” refers to a newly sequenced set of 73 traditional Chinese varieties (**table S1**). (**B**) Fit of observed f4 statistics with the graph. “Balsas” refers to the Balsas samples of *parviglumis.* Observed estimates of f4 are shown as short vertical lines with error bars (black lines) while circles show the expected values generated by the graph that are within (green) or outside of (red) the range observed in the data.

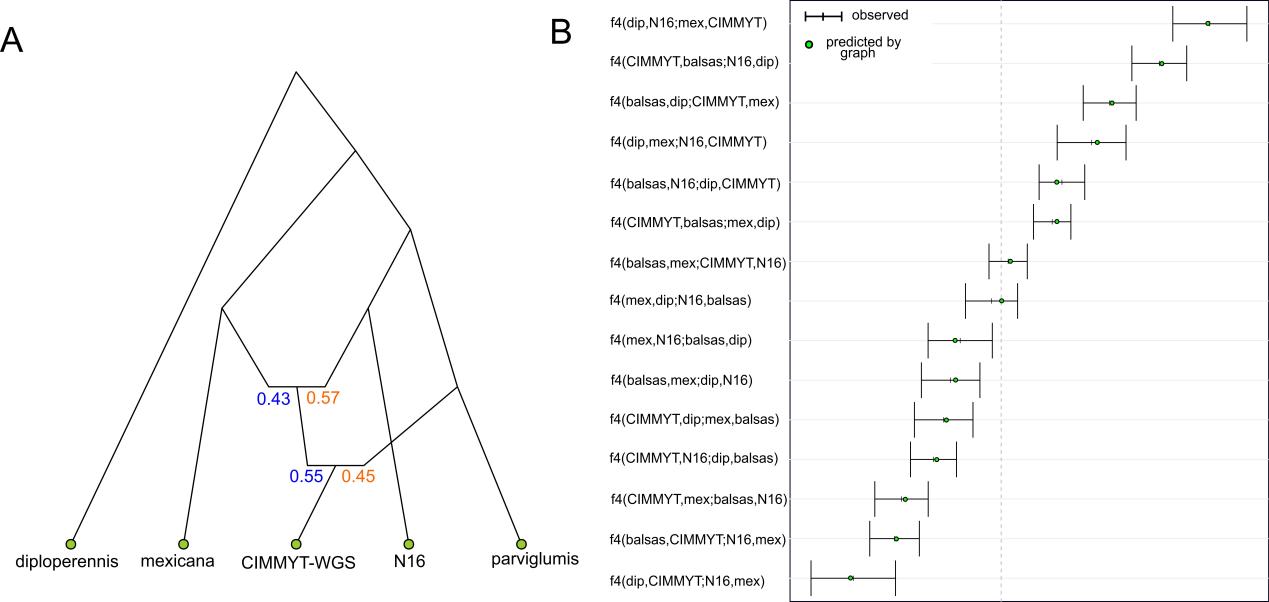

**Figure S9** (**A**) Admixture graph of landraces (CIMMYT-WGS) and teosinte. Colors in blue and yellow at each node represent the estimated proportion of admixture from each source population. “CIMMYT-WGS” refers to a newly sequenced 267 open-pollinated traditional maize varieties from across Mexico (**table S1**). (**B**) Fit of observed f4 statistics with the graph. Observed estimates of f4 are shown as short vertical lines with error bars (black lines) while circles show the expected values generated by the graph that are within (green) or outside of (red) the range observed in the data.

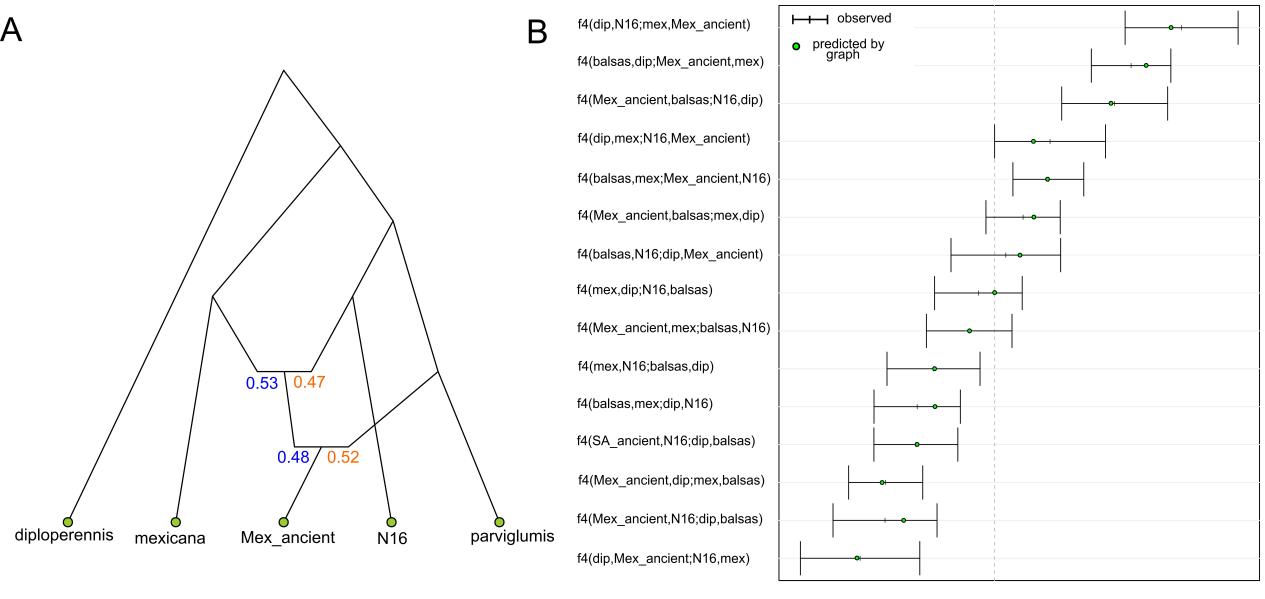

**Figure S10** (**A**) Admixture graph of ancient maize (Mex_ancient) and teosinte. Colors in blue and yellow at each node represent the estimated proportion of admixture from each source population. “Mex_ancient” refers to the 5,310-year-old maize-Tehuacan162 (**table S1**) (*41*). (**B**) Fit of observed f4 statistics with the graph. Observed estimates of f4 are shown as short vertical lines with error bars (black lines) while circles show the expected values generated by the graph that are within (green) or outside of (red) the range observed in the data.

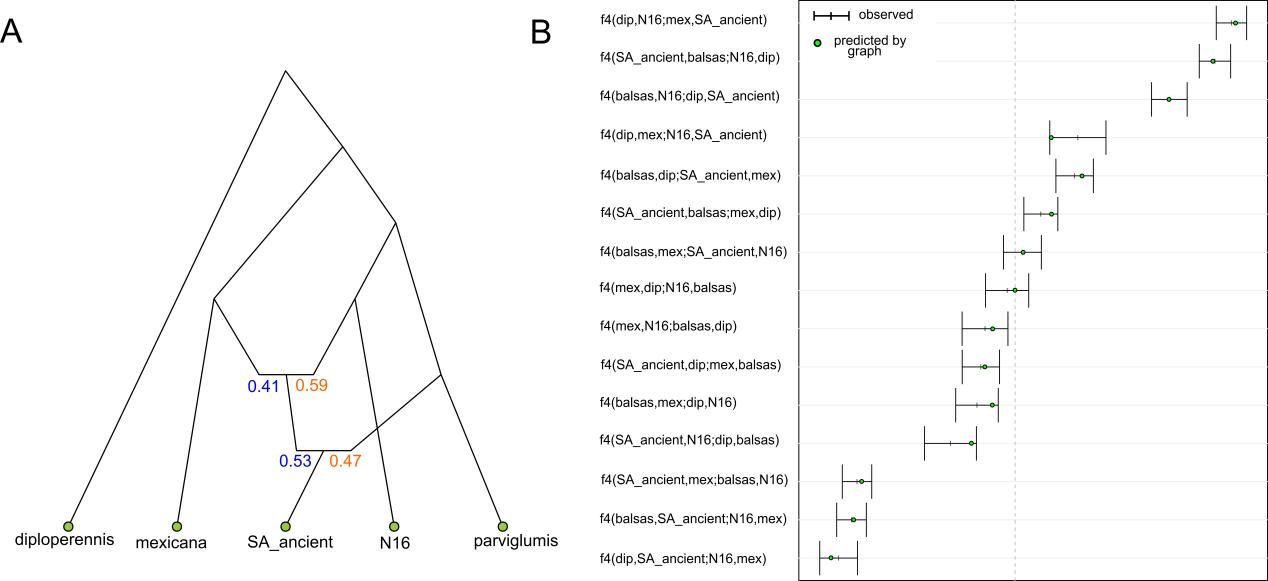

**Figure S11** (**A**) Admixture graph of ancient maize (SA_ancient) and teosinte. Colors in blue and yellow at each node represent the estimated proportion of admixture from each source population. “SA_ancient” refers to the ancient maize from South America dating to ~1,000 cal BP including 7 samples(**table S1**) (*42*). (**B**) Fit of observed f4 statistics with the graph. Observed estimates of f4 are shown as short vertical lines with error bars (black lines) while circles show the expected values generated by the graph that are within (green) or outside of (red) the range observed in the data.

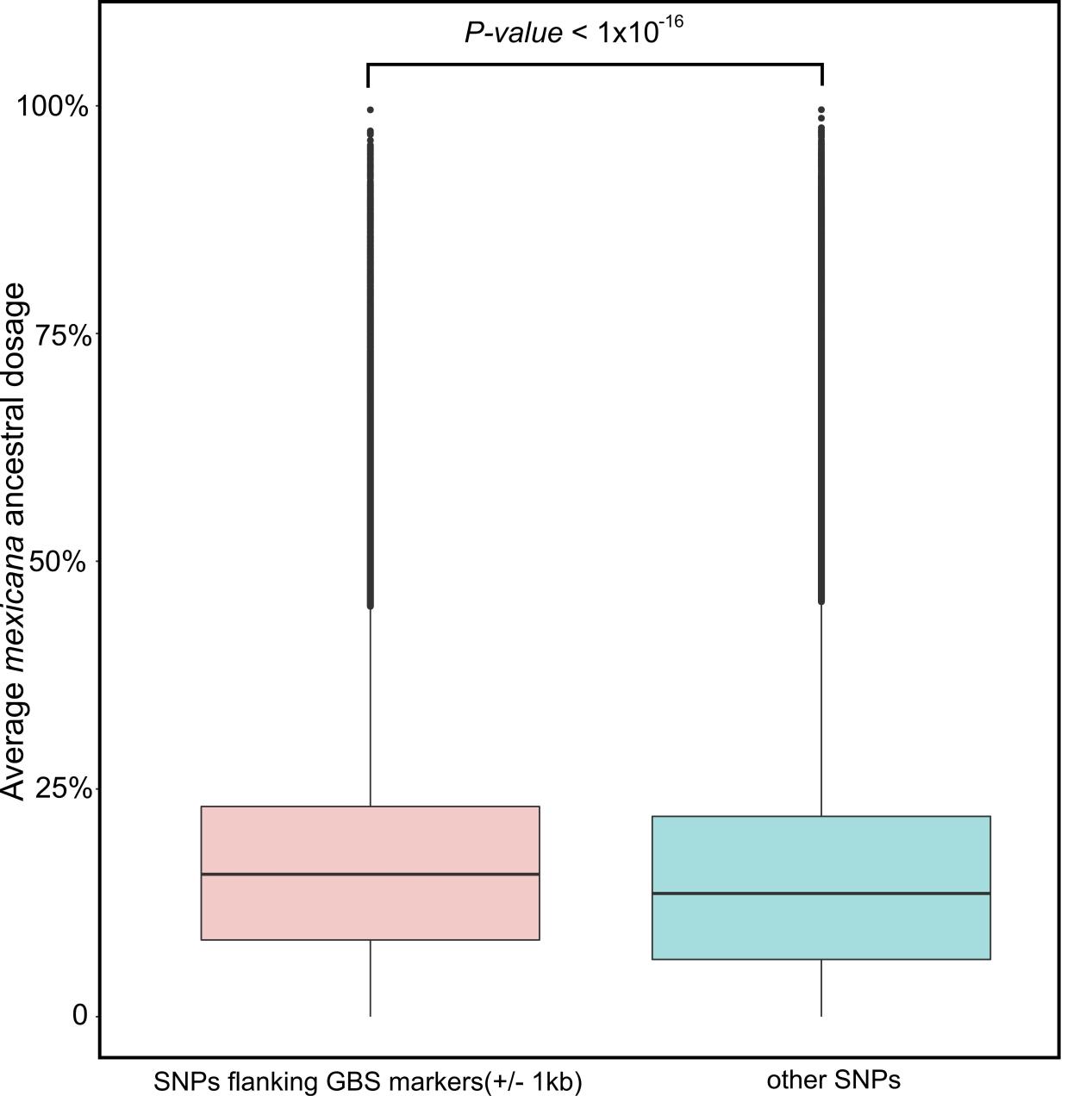

**Fig. S12.** **Percent *mexicana* ancestry comparison of SNPs flanking GBS markers and SNPs far from GBS markers.** Boxplot of average mexicana ancestral dosage estimated by ELAI for SNPs of 507 maize within 1kb of a GBS marker (red) and those more distant (blue). The *P* value was determined by the Kolmogorov-Smirnov test.

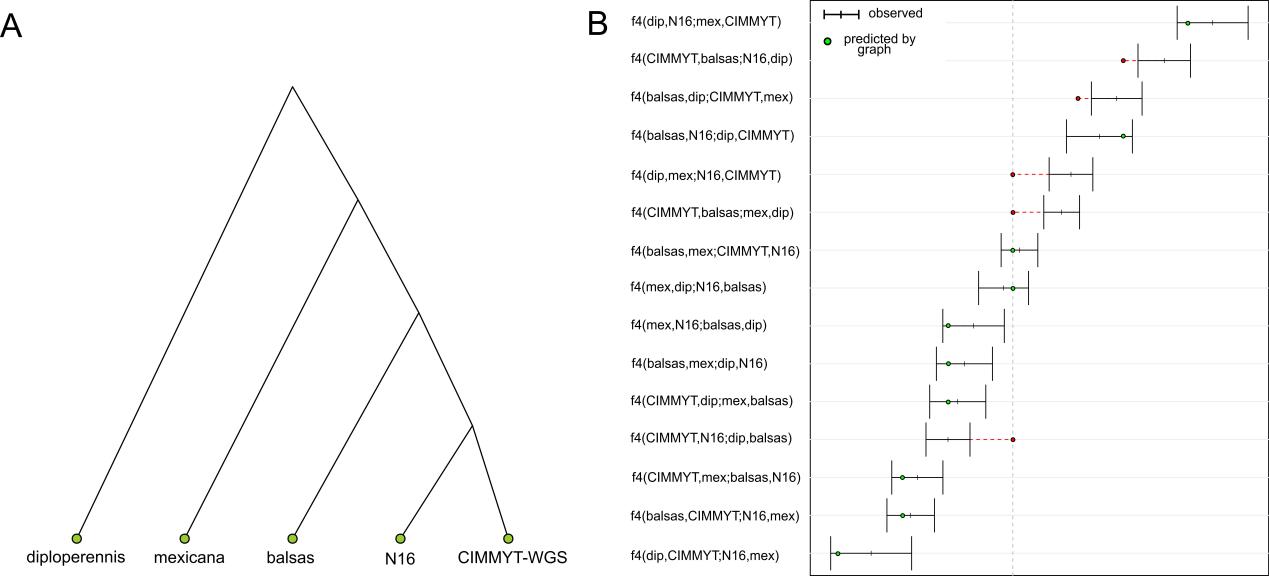

**Fig. S13.** (**A**) “No admixture” admixture graph of landraces (CIMMYT-WGS) and teosinte. “CIMMYT-WGS” refers to a newly sequenced 267 open-pollinated traditional maize varieties from across Mexico (**table S1**). In this graph, we assumed there was no subsequent admixture between landraces from across Mexico and *parviglumis*/*mexicana* after the domestication of maize. (**B**) Fit of observed f4 statistics with the graph. “Balsas” refers to the Balsas samples of *parviglumis.* Observed estimates of f4 are shown as short vertical lines with error bars (black lines) while circles show the expected values generated by the graph that are within (green) or outside of (red) the range observed in the data.

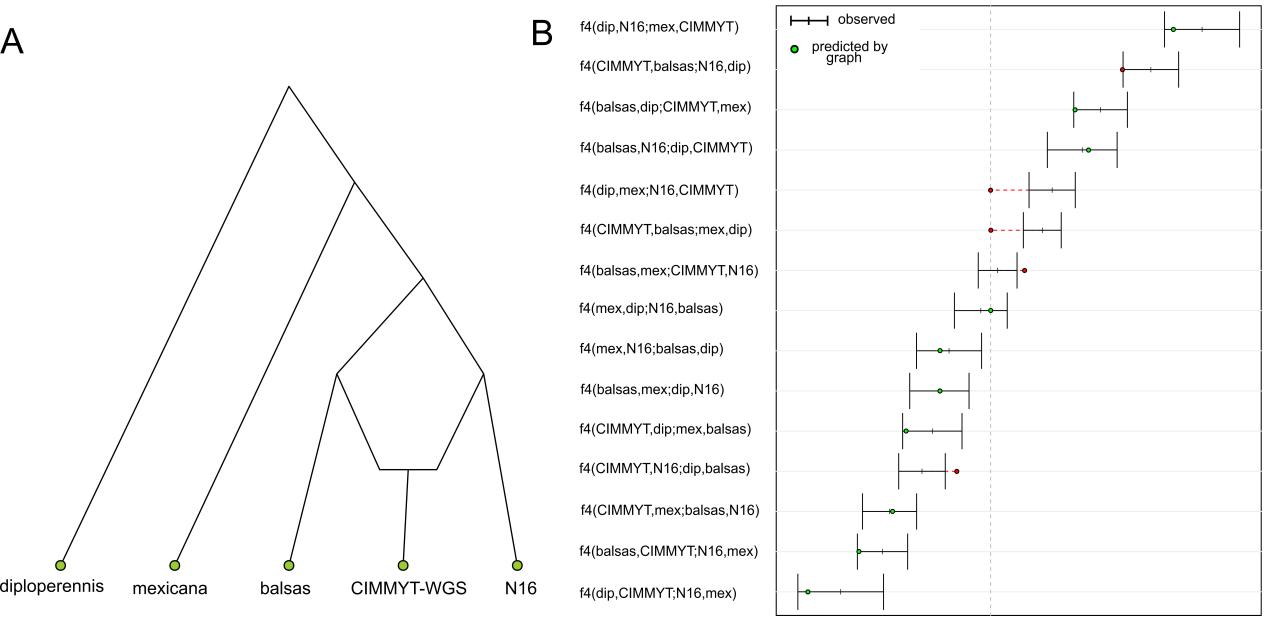

**Fig. S14.** (**A**) “One time admixture” admixture graph of landraces (CIMMYT-WGS) and teosinte. “CIMMYT-WGS” refers to a newly sequenced 267 open-pollinated traditional maize varieties from across Mexico (**table S1**). In this graph, we assumed there was only one subsequent admixture between landraces from across Mexico and *parviglumis* after the domestication of maize. (**B**) Fit of observed f4 statistics with the graph. “Balsas” refers to the Balsas samples of *parviglumis.* Observed estimates of f4 are shown as short vertical lines with error bars (black lines) while circles show the expected values generated by the graph that are within (green) or outside of (red) the range observed in the data.

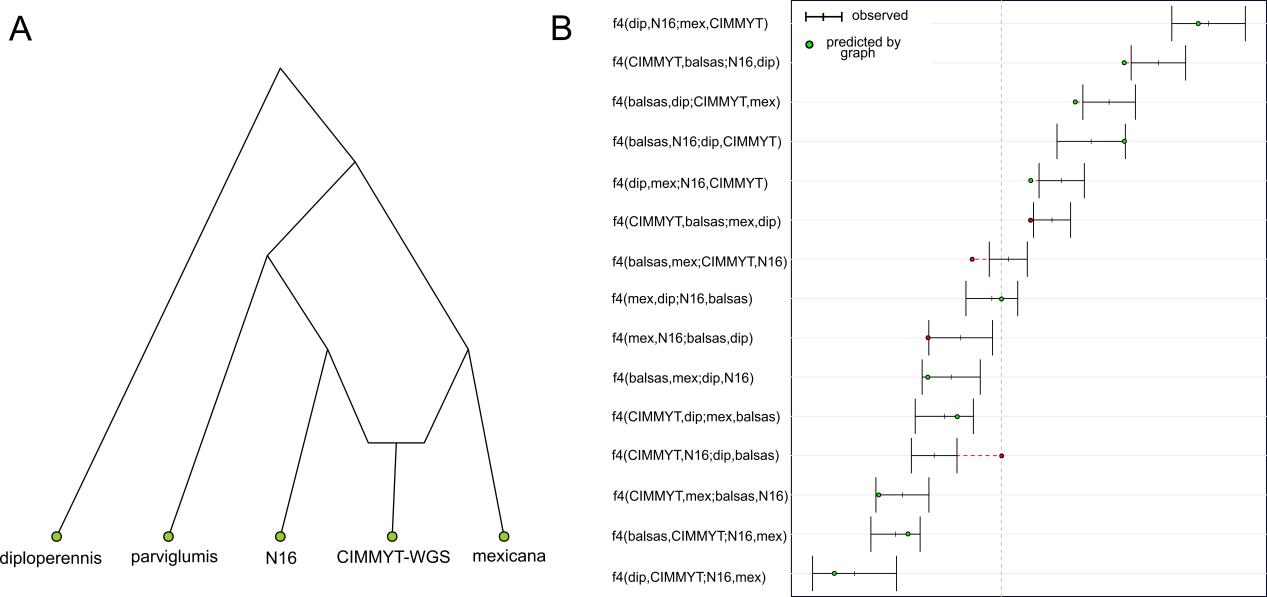

**Fig. S15.** (**A**) The other “One time admixture” admixture graph of landraces (CIMMYT-WGA) and teosinte. “CIMMYT-WGS” refers to a newly sequenced 267 open-pollinated traditional maize varieties from across Mexico (**table S1**). In this graph, we assumed there was only one subsequent admixture between landraces from across Mexico and *mexicana* after the domestication of maize. (**B**) Fit of observed f4 statistics with the graph. “Balsas” refers to the Balsas samples of *parviglumis.* Observed estimates of f4 are shown as short vertical lines with error bars (black lines) while circles show the expected values generated by the graph that are within (green) or outside of (red) the range observed in the data.

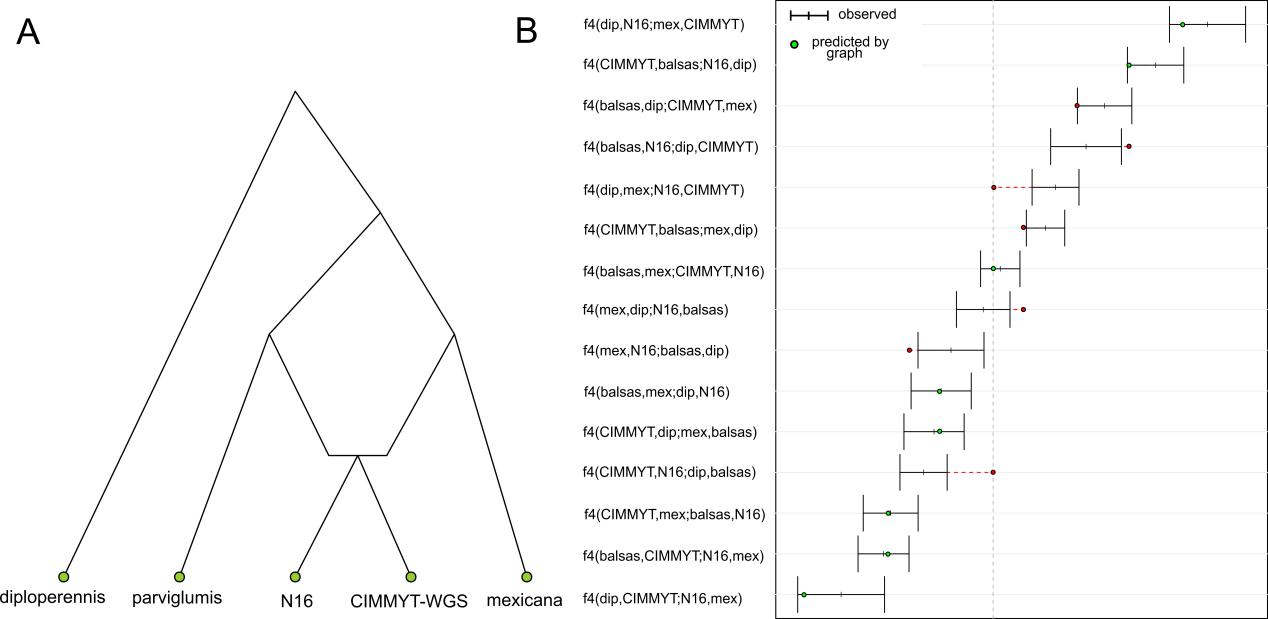

**Fig. S16.** (**A**) Admixture graph of landraces (CIMMYT-WGS) and teosinte. “CIMMYT-WGS” refers to a newly sequenced 267 open-pollinated traditional maize varieties from across Mexico (**table S1**). In this graph, we assumed that N16 and the landraces from across Mexico all derived from the hybrid between *parviglumis* and *mexicana.* (**B**) Fit of observed f4 statistics with the graph. “Balsas” refers to the Balsas samples of *parviglumis.* Observed estimates of f4 are shown as short vertical lines with error bars (black lines) while circles show the expected values generated by the graph that are within (green) or outside of (red) the range observed in the data.

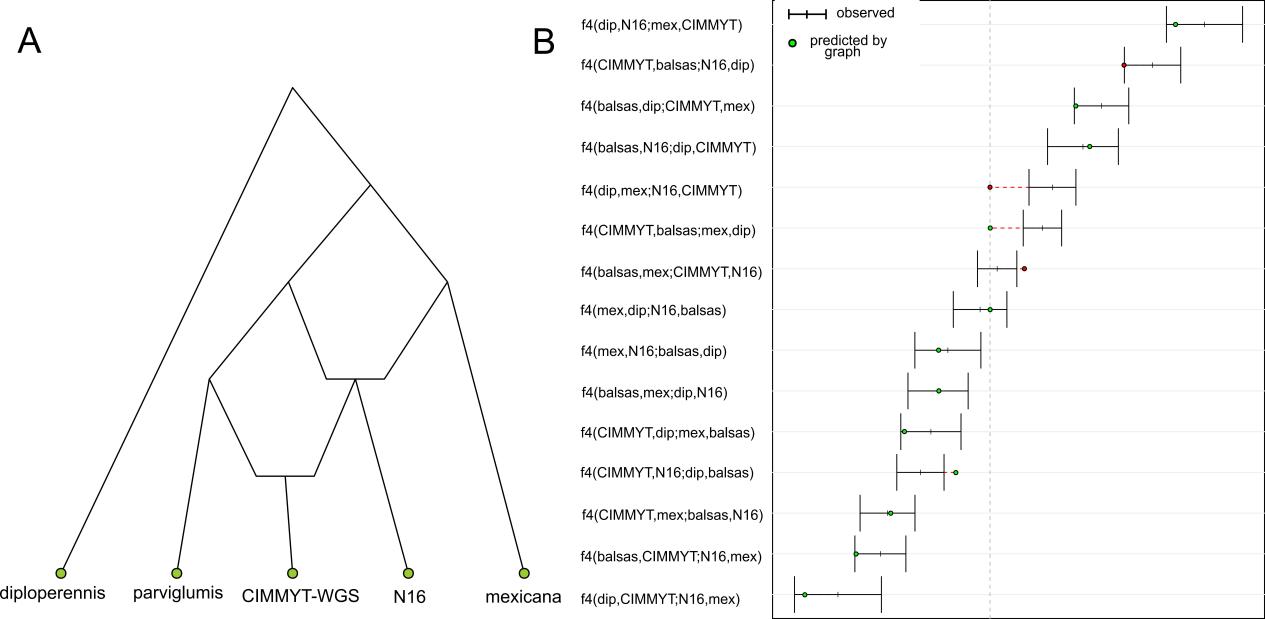

**Fig. S17.** (**A**) Admixture graph of landraces (CIMMYT-WGS) and teosinte. “CIMMYT-WGS” refers to a newly sequenced 267 open-pollinated traditional maize varieties from across Mexico (**table S1**). In this graph, we assumed that N16 derived from the hybrid between *parviglumis* and *mexicana*, however, landraces from across Mexico derived from the second hybridization event between the hybrid from the first hybridization event and *parviglumis.* (**B**) Fit of observed f4 statistics with the graph. “Balsas” refers to the Balsas samples of *parviglumis.* Observed estimates of f4 are shown as short vertical lines with error bars (black lines) while circles show the expected values generated by the graph that are within (green) or outside of (red) the range observed in the data.

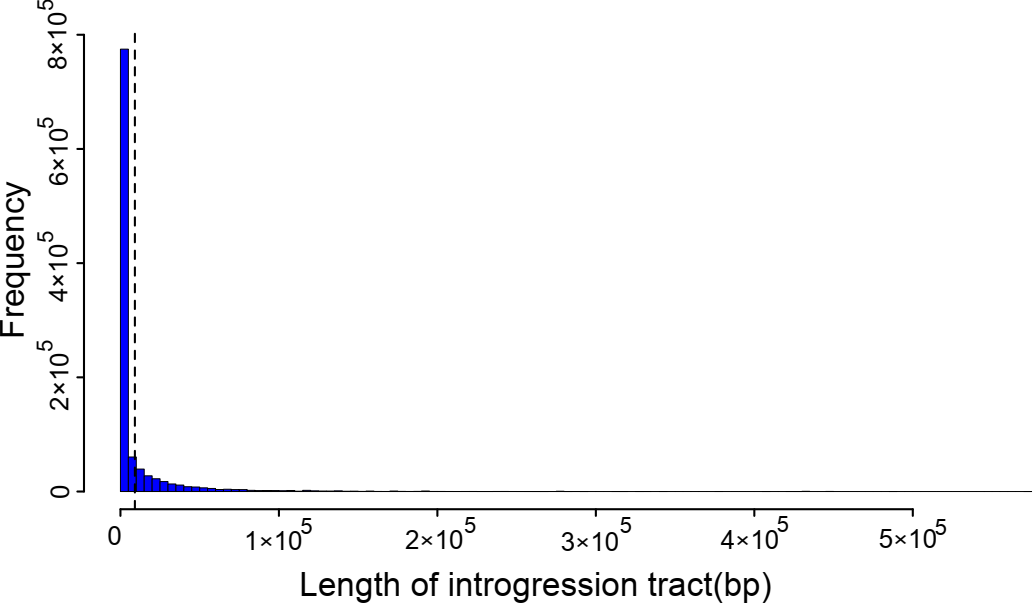

**Fig. S18. Distribution of the length of *mexicana* introgression tracts in 845 maize estimated using ELAI.**

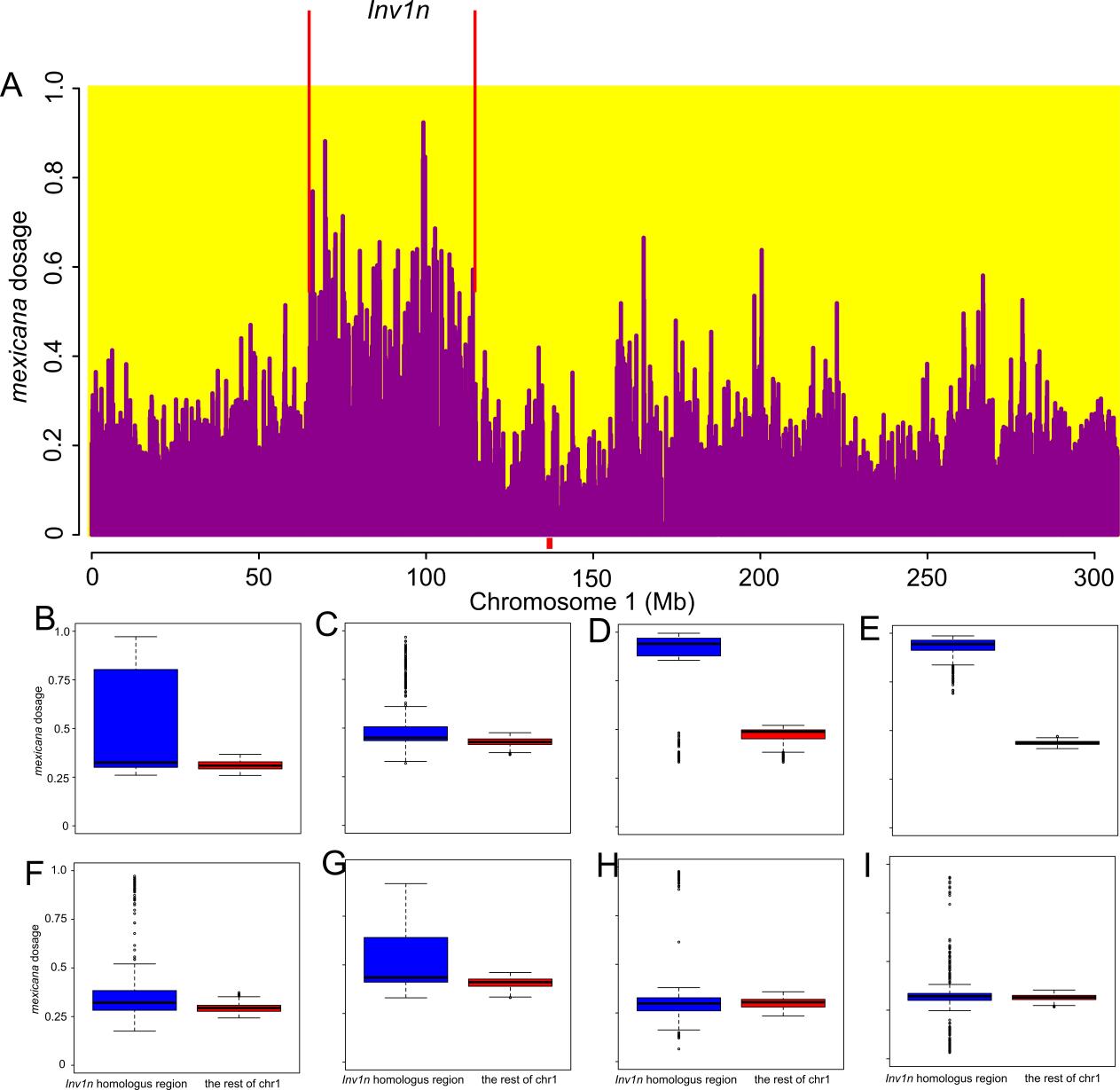

**Fig. S19. The average *mexicana* dosage of 845 maize inbreds and landraces across chromosome 1.** (**A**) There is a clear elevation of mexicana dosage in the *Inv1n*. (**B**-**I**) *mexicana* dosage estimated by ELAI in *Inv1n* regions using *parviglumis* reference samples with *Inv1n* (blue) and *parviglumis* without *Inv1n* as reference populations (red). The inversion genotypes at these 8 regions within *Inv1n* were from Chen *et al*. (*39*).

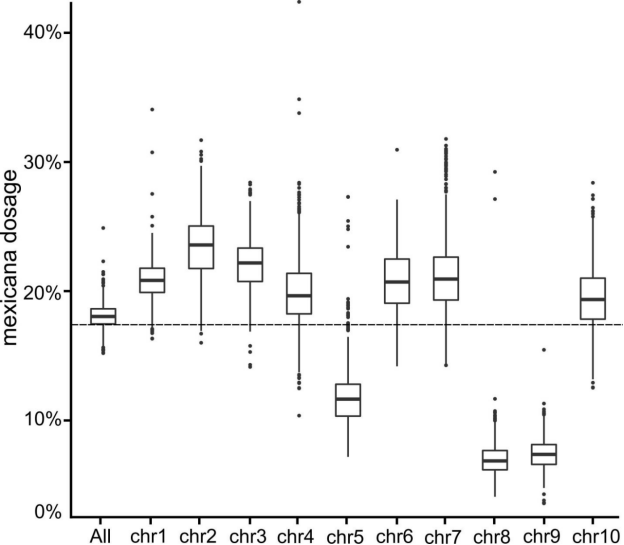

**Fig. S20. *mexicana* dosage of each chromosome.** The dashed line indicates the mean *mexicana* dosage of the whole genome. Boxes show the median and interquartile range for each chromosome, and whiskers represent 1.5 s.d. of the interquartile range.

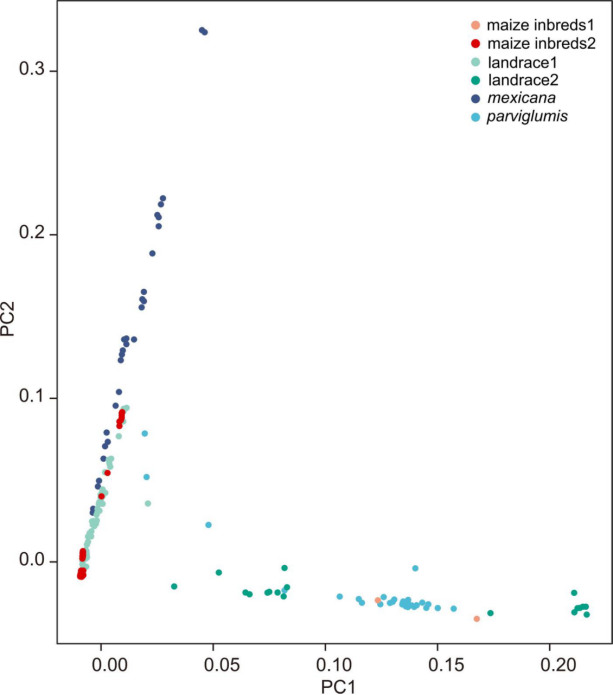

**Fig. S21.** **Principal component analysis (PCA) of *ZmPRR37a*** (±2Kb) **for 845 maize, *mexicana* and *parviglumis*.** Each point represents one accession. Only few modern maize inbred lines(maize in breds1) and part of landraces (landrace2) cluster with *parviglumis*, while most modern maize inbred lines (maize inbreds2) and landraces(landrace1) clusters with *mexicana* haplotypes.

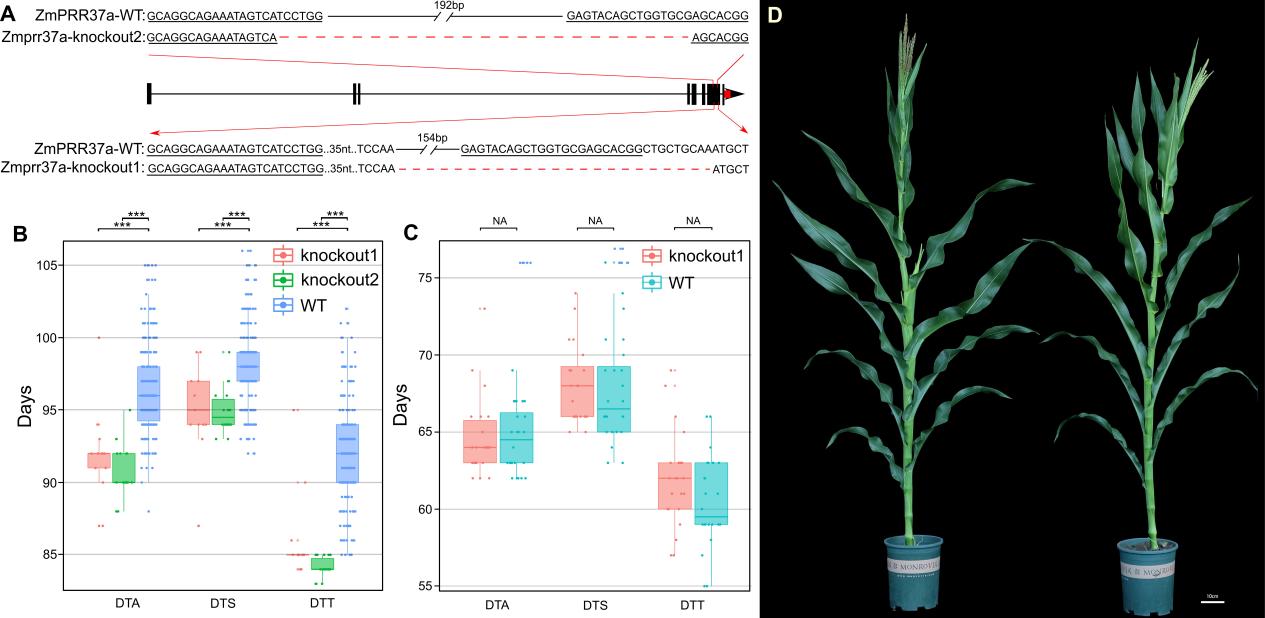

**Fig. S22. Phenotype analysis of different CRISPR/Cas9 mutation *ZmPRR37a* in different environments.** (**A**) Gene structure and sequences of *ZmPRR37a* target regions in wild type, *ZmPRR37a* CRISPR/Cas9 knockout mutants 1 and 2; (**B**) Statics of days to anthesis, days to silk, days to tassel in Jilin province (one long day condition, 2018; China E124°49’, N43°30’) of knockout1 (n=13), knockout2 (n=18) and wild type (n=234); (**C**) Days to anthesis, days to silk, days to tassel in Hainan province (one short day condition,2020; China; E108°43’, N18°34’) of knockout1 (n=22) and wild type (n=24); (**D**) *ZmPRR37a* knockout (left) and WT (right) maize grown in long day conditions. Scale bar 10cm.

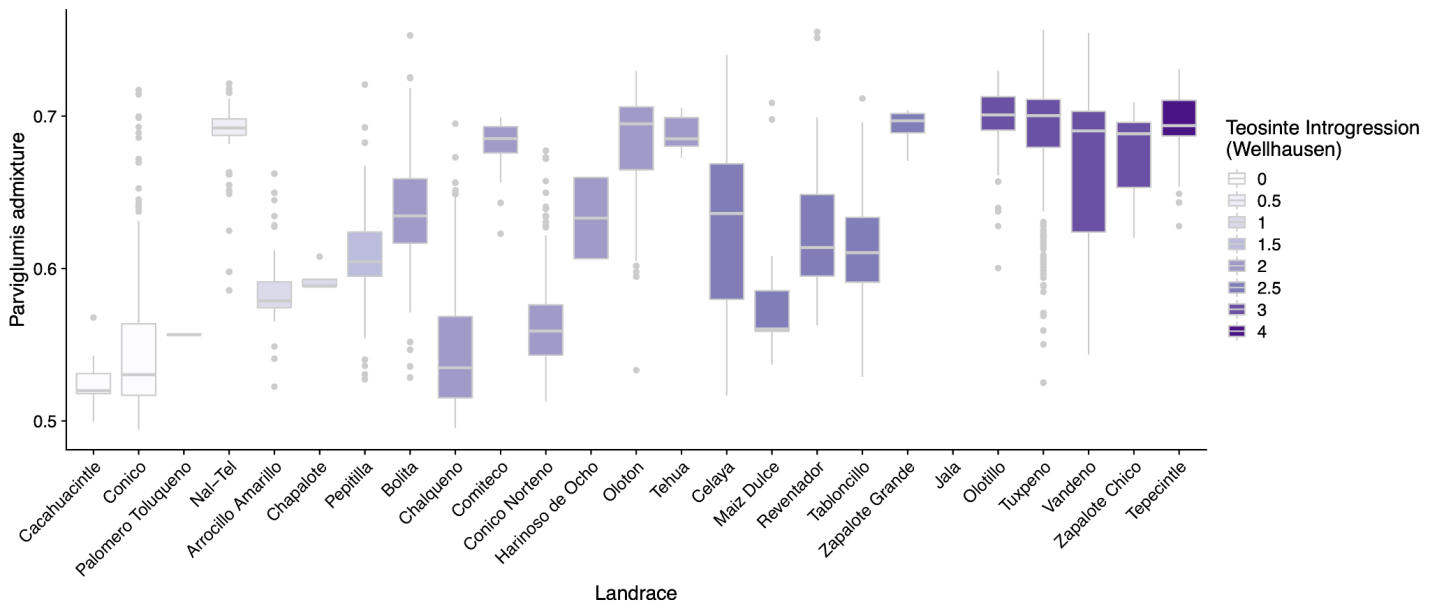

**Fig. S23. Historical landrace teosinte introgression is related to STRUCTURE estimates of parviglumis ancestry**. Historical values summarizing ear morphology range from 0=no teosinte introgression to 4=very strong teosinte introgression, and landraces are ranked along the x-axis by increasing teosinte introgression. The Spearman correlation between teosinte introgression and parviglumis ancestry is positive, statistically significant, and very large (ρ = 0.67, S = 2.67e+08, p < .001).

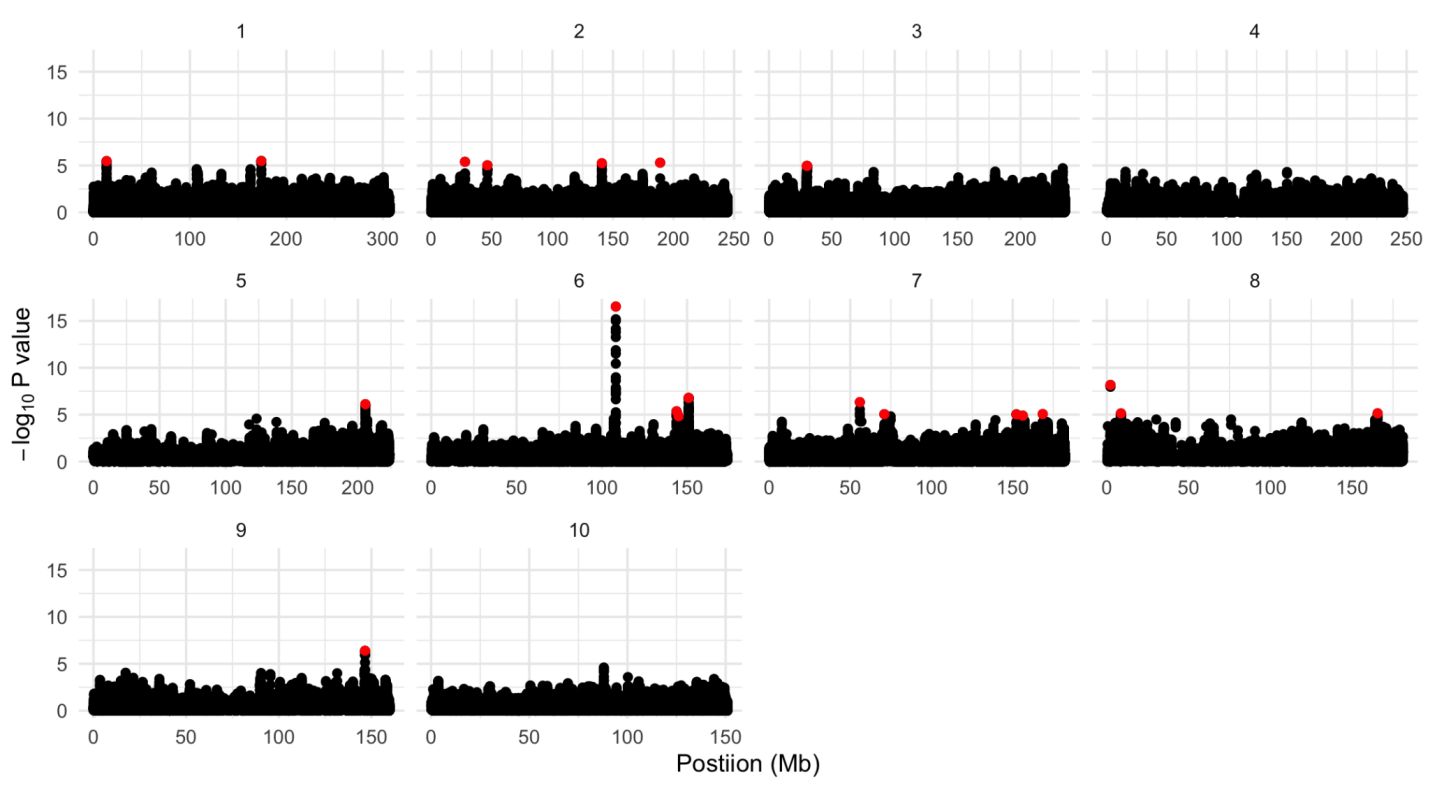

**Fig. S24. Manhattan plot of multivariate admixture association mapping in maize inbred lines.** Y-axis represents the F-test p-value. Red points highlight SNPs significant at a false discovery rate of 10%

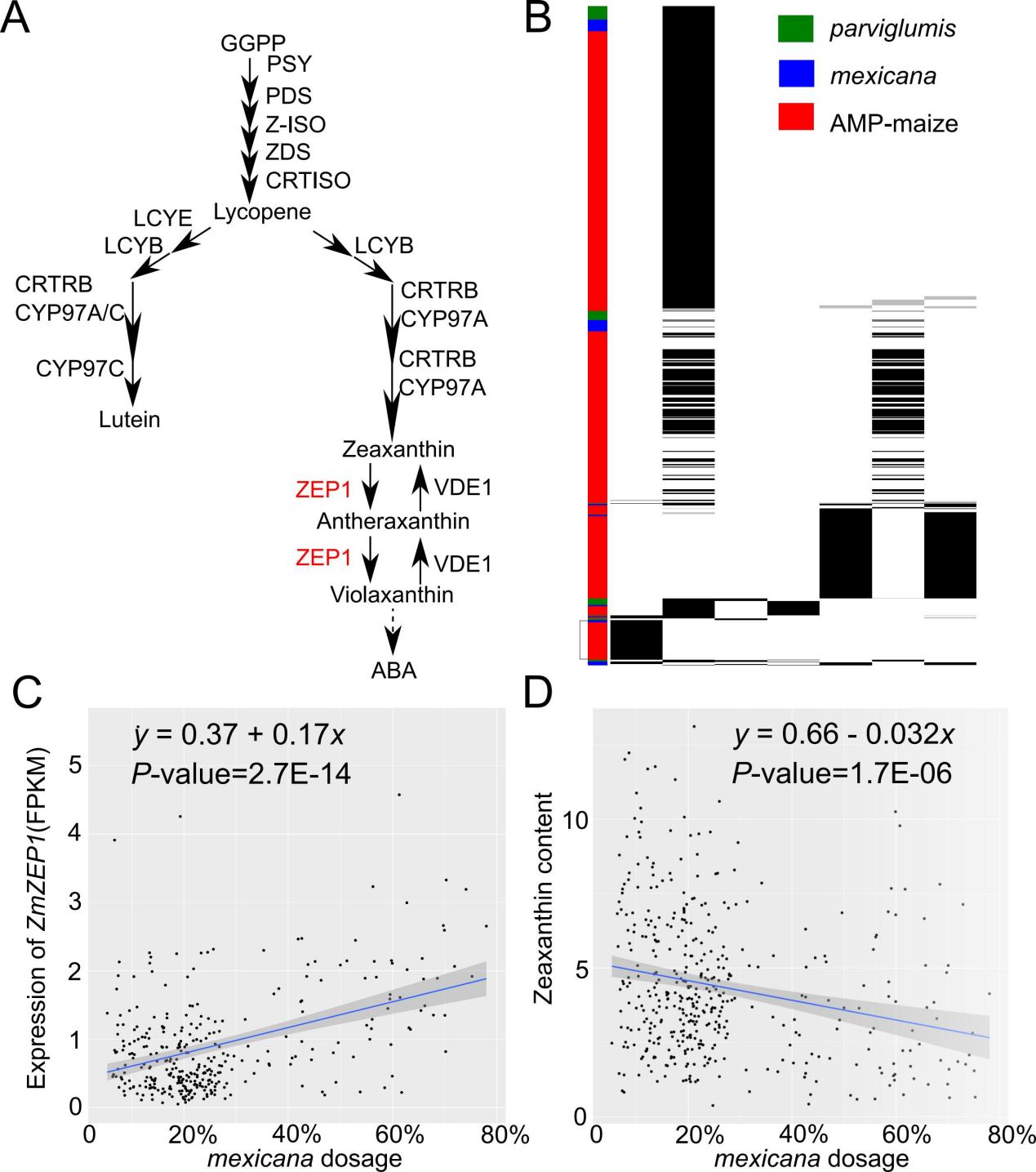

**Fig. S25. The *mexicana* haplotype increases gene expression of *ZmZEP1* and reduces zeaxanthin content in maize kernels.** (**A**) Metabolic pathway showing the role of *ZmZEP1* in modulating *Zeaxanthin* content. (**B)** Haplotstrip (*125*) diagram of maize, *parviglumis*, and *mexicana* at *ZmZEP1.* (**C)** *mexicana* dosage estimated by ELAI positively correlates with *ZmZEP1* expression. (**D**) *mexicana* dosage negatively correlates with zeaxanthin content in kernels.

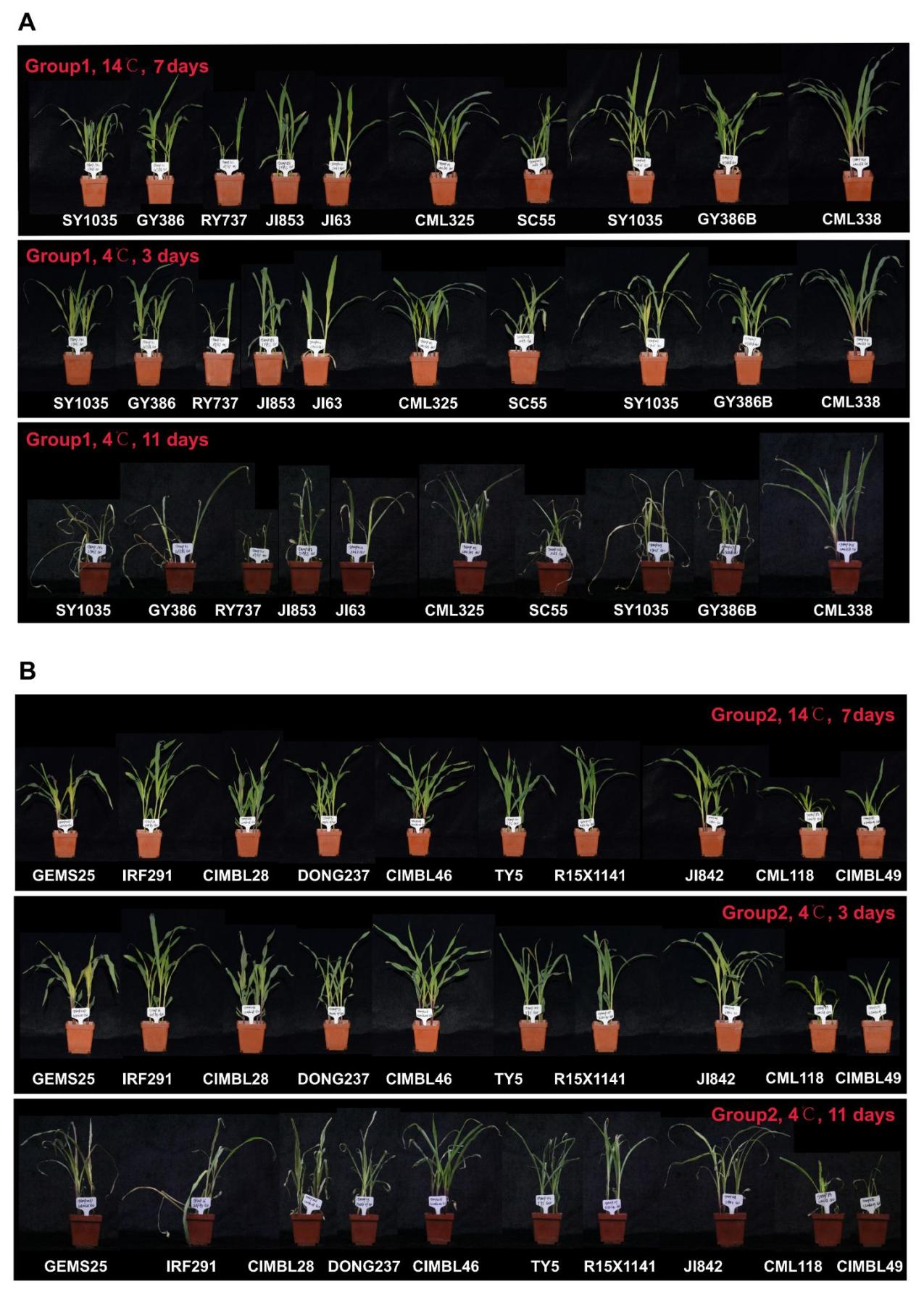

**Fig. S26. High mexicana dosage maize and low mexicana dosage maize at dgat1 locus didn't display obvious cold tolerance differences.** Chilling experiments exposed plants to cold acclimation (14℃,7 days) and then 3 days at 4 ℃ followed by 3 days recovery at normal conditions (28 ℃) (**A**) Images of 10 inbreds of high *mexicana* dosage at *dgat1,* (**B**) Images of 10 inbreds of low *mexicana* dosage at *dgat1.*

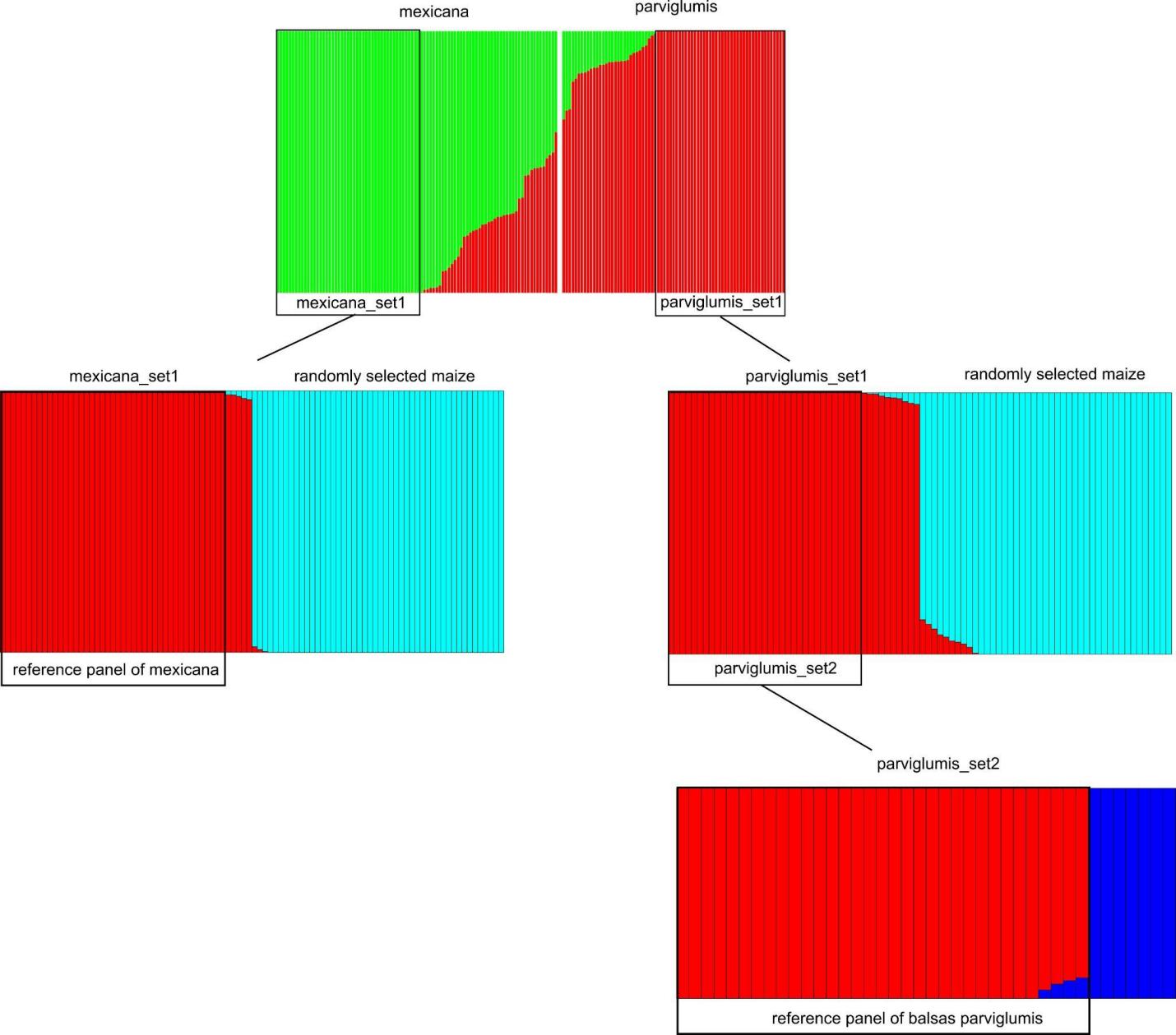

**Fig. S27. Reference panel of *parviglumis* and *mexicana*.** Results of ADMIXTURE (*103*) run using 1,665,173 SNPs from *mexicana* and *parviglumis* individuals from Chen *et al*. (*39*). From the initial ADMIXTURE of those two taxa with k=2 (top), we chose unadmixed individuals to compare in a second model with k=2 to 48 maize for *mexicana* and 43 maize for *parviglumis* groups separately (middle). Individuals showing no admixture with maize were then retained. *parviglumis* was further subdivided given the deep split often seen between populations in the Balsas and Jalisco regions (*22*) (bottom).

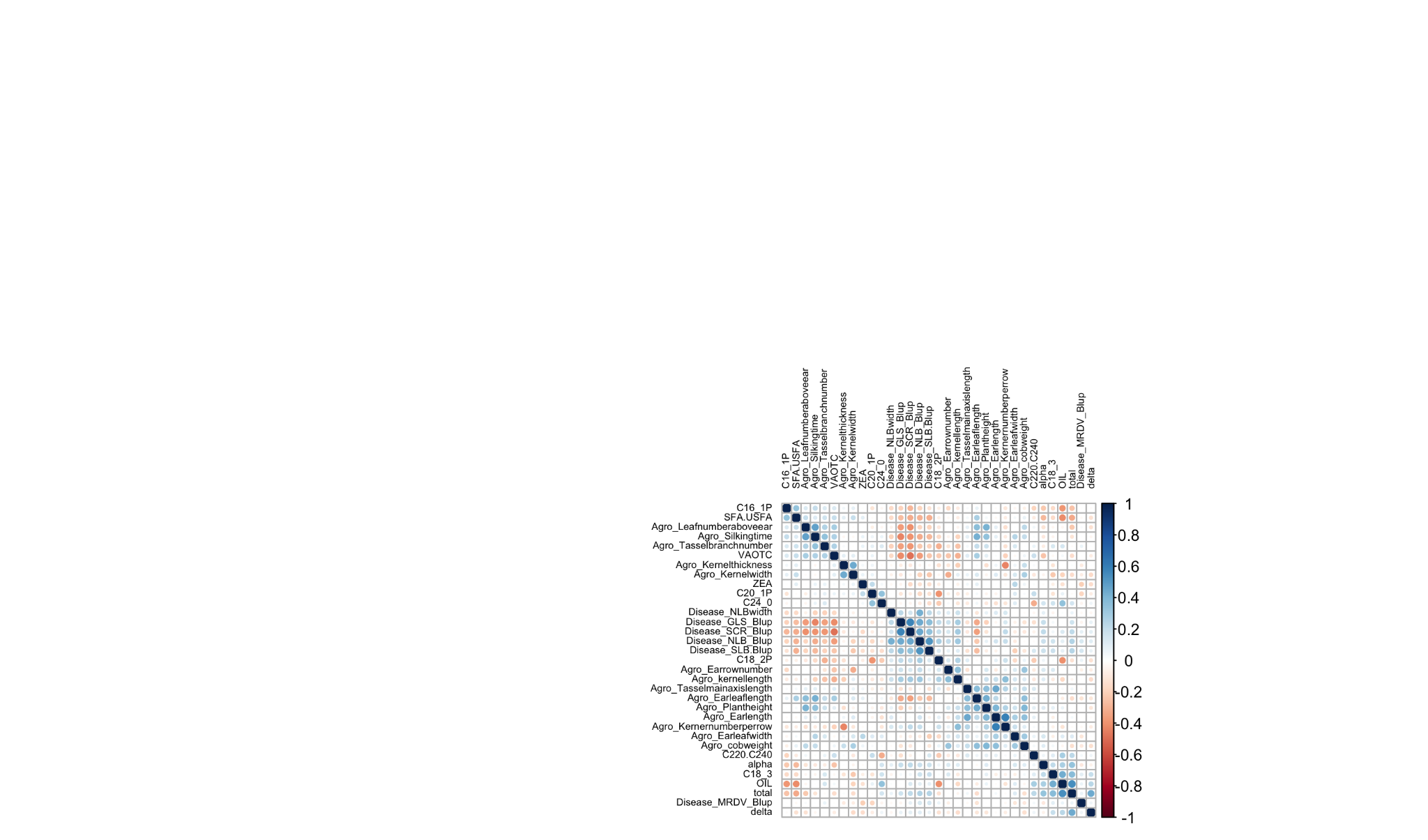

**Fig. S29.** Correlation matrix of log-scaled phenotypes for 33 traits.

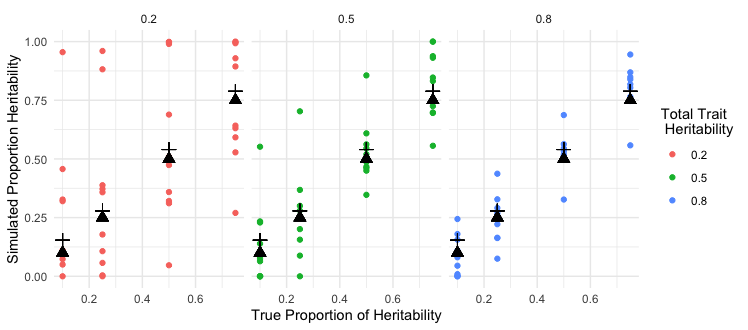

**Fig. S30. Simulated trait heritability:** Ten replicates each of twelve sets of simulated heritabilities were generated using the empirical kinships partitioned on introgressed regions and the entire genome. Colors represent the total trait h^2^, means of the 10 replicates are shown as a black cross, and the true proportion of h^2^ due to admixture is shown as a black triangle.
